# A Fragment Screen Identifies Acrylamide Covalent Inhibitors of the TEAD•YAP Protein-Protein Interaction

**DOI:** 10.64898/2026.03.18.712694

**Authors:** Khuchtumur Bum-Erdene, Mona K. Ghozayel, Mark J. Zhang, Giovanni Gonzalez-Gutierrez, Samy O. Meroueh

## Abstract

TEA domain (TEAD) proteins bind co-activator Yes-associated protein (YAP) to regulate the expression of target genes of the Hippo pathway. The TEAD•YAP protein-protein interaction is not druggable, but TEADs possess a unique and deep palmitate pocket with a highly conserved cysteine located outside the TEAD•YAP protein-protein interaction interface. Here, we screen a fragment library of acrylamide electrophiles and identify a fragment that forms an adduct with the conserved palmitate pocket cysteine and inhibits TEAD4 binding to YAP. Synthesis of a focused set of derivatives and time- and concentration-dependent studies with four TEADs provide reaction rates and binding constants. Co-crystal structures of fragments bound to TEAD2 and TEAD3 reveal reaction at the conserved palmitate pocket cysteine but also at another less conserved cysteine located in the palmitate pocket of TEAD2 closer to the TEAD•YAP interface. These fragments provide a starting point for the development of allosteric acrylamide small-molecule covalent TEAD•YAP inhibitors.

## INTRODUCTION

TEA domain (TEAD) transcription factors engage the yes-associated protein (YAP) or its paralog transcriptional coactivator with PDZ-binding motif (TAZ) to create a protein-protein interaction complex that controls transcriptional activity of the Hippo pathway. In normal tissue, YAP and TAZ are phosphorylated by Hippo kinases Large tumor suppressor-1/2 (Lats1/2) and sequestered in the cytoplasm by 14-3-3 proteins (1). Cytoplasmic YAP and TAZ are also degraded by the proteasome (2, 3). In cancer, loss of cell contacts with other cells or with the extracellular matrix turns off Hippo. As a result, YAP and TAZ are no longer phosphorylated, so they are translocated to the nucleus to bind to TEADs and other transcription factors. The TEAD•YAP/TAZ complex target genes include *CTGF*, *CYR61*, *MYC*, *WNT5A/B*, *EGFR* and *PD-L1* that are associated with tumor growth and metastasis (4–16).

The TEAD family includes four paralogs, TEADs1-4. Each TEAD possesses conserved DNA-and YAP- or TAZ-binding domains. The binding affinity of TEADs to YAP is high (17, 18), and the binding interface is more than 1000 Å^2^ (19, 20). The interface lacks a well-defined binding pocket for small-molecule binding, so it is considered undruggable. However, TEADs possess a unique deep and well-defined palmitate-binding pocket. The palmitate lipid is completely buried in the pocket, and it forms a thioester bond with a conserved cysteine residue at the entrance of the palmitate pocket (21, 22). The palmitate does not alter TEAD binding to YAP (17, 18). TEAD stability and cellular half-life are enhanced by the binding of palmitate (18, 21–23). Previously, we reported a strategy that led to inhibition of the TEAD•YAP protein-protein interaction based on the approved drug flufenamic acid (17). We designed a small molecule, TED-347, which formed a covalent bond with conserved cysteine of the palmitate pocket and inhibited TEAD binding to YAP in a concentration- and time-dependent manner (17). Since then, a number of covalent (24–33) and non-covalent (34–40) compounds have been reported that bind to this pocket. Genentech researchers reported a non-covalent small molecule that binds to the palmitate pocket and inhibits TEAD binding to YAP (39). We recently reported small-molecule inhibitors of TEAD binding to YAP using a cyanamide warhead (24).

Here, we screen a library of acrylamide electrophile fragments to identify small molecules that compete with palmitate and inhibit TEAD binding to YAP. We used a fluorescence polarization (FP) assay for the screening. Hit fragments were further characterized by whole protein mass spectrometry to confirm adduct formation at the palmitate pocket cysteine. We selected one fragment, **1** (ACR-021), and designed and synthesized derivatives that led to small-molecule covalent inhibitors of TEAD4 binding to YAP. Time- and concentration-dependent studies with all four TEADs and for all compounds provided second order rate constants and revealed selectivity of these compounds for TEAD1 and TEAD3. Co-crystal structures with TEAD2 and TEAD3 revealed the binding mode of these compounds.

## RESULTS

### Acrylamide Fragment Screen

A library of 372 acrylamide fragments was screened for inhibition of TEAD4 binding to YAP using a fluorescence polarization assay that uses a fluorescently labeled TEAD-binding domain of YAP (FAM-YAP_60–99_) (17). TEAD4 was incubated with acrylamide fragments (50 µM) for 24 h at 4°C and binding was detected by fluorescence polarization (**Fig. 1A**). Thirteen fragment hits inhibited TEAD4•YAP binding by more than 40%. Except for fragments **1** (ACR-021) and **2** (ACR-362) [**Fig. 1B**], fragments **3**-**13** (**Fig. S1**) inhibited both wildtype TEAD4 and TEAD4^C367S^ with similar double-digit micromolar IC_50_s (**Table S1, Fig. S2A,** and **S2B**). These compounds inhibit mutant TEAD4^C367S^ binding to YAP. As a result, the inhibition of the TEAD4•YAP interaction by these fragments is unlikely due to covalent bond formation at the central pocket. We do not pursue these further, as the focus of this manuscript is only on covalent inhibitors of the TEAD•YAP protein-protein interaction.

**Figure 1.**
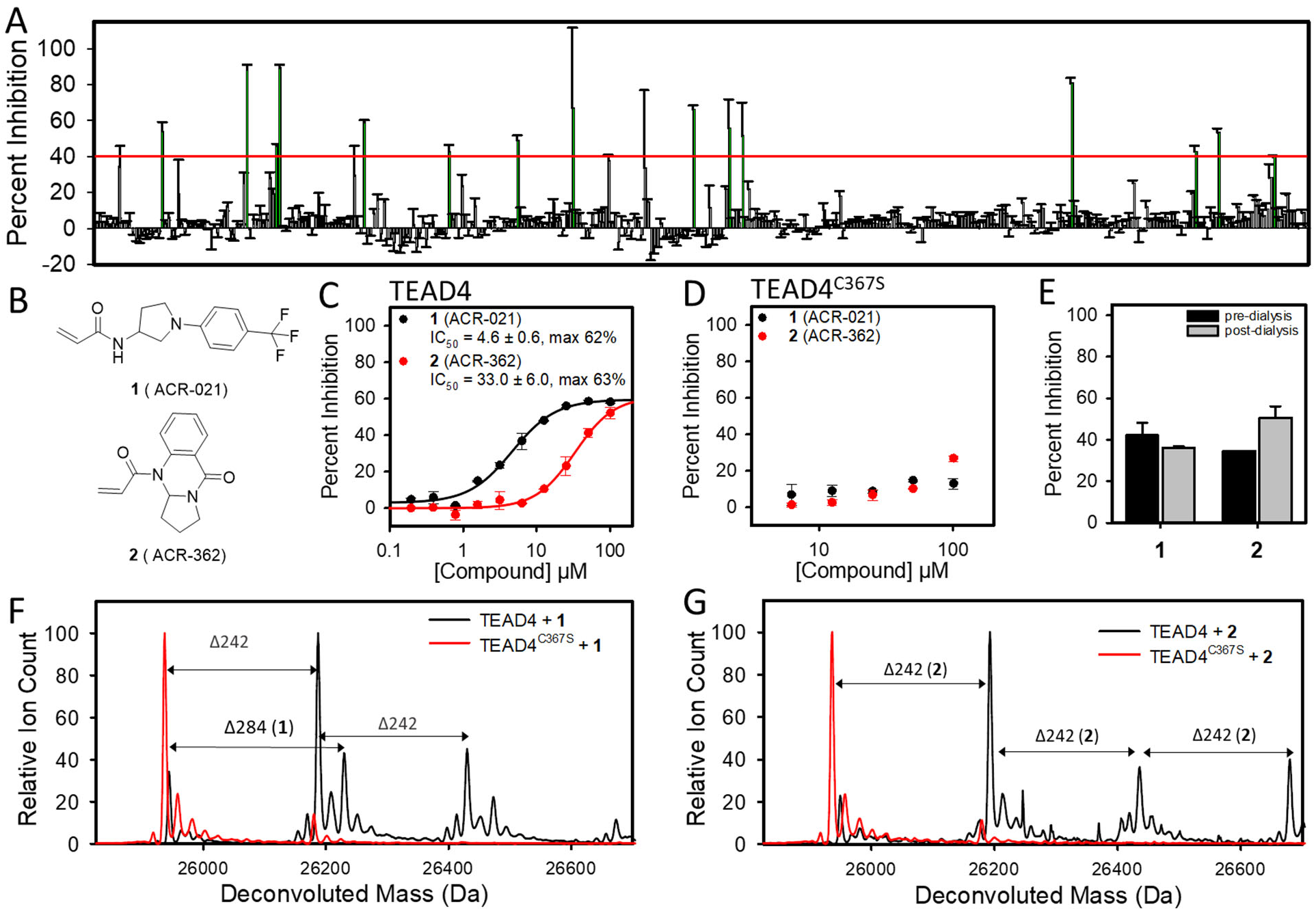
Acrylamide Fragment Screen Hits**. (A)** TEAD4 was preincubated with 50 µM acrylamide fragments for 24 h at 4°C and tested for binding to fluorescently labeled YAP (residues 60-99) peptide (n = 2). **(B)** Chemical structures of fragments **1** (ACR-021) and **2** (ACR-362). **(C)** TEAD4 was preincubated with 0.2-100 µM fragments for 24 h at 4°C before measuring its binding to the fluorescently labeled YAP peptide. Percent inhibition was determined relative to DMSO control and a sample without TEAD4 (mean ± s.d., n = 3). **(D)** TEAD4^C367S^ was preincubated with 6.25-100 µM fragments for 24 h at 4°C before measuring its binding to the fluorescently labeled YAP peptide. Percent inhibition was determined relative to DMSO control and a sample without TEAD4 (mean ± s.d., n=3). **(E)** TEAD4 was preincubated with 100 µM compounds for 24 h at 4°C and dialyzed against buffer for 24 h at 4°C. Detection of binding to YAP peptide was performed before and after dialysis of the samples (n = 2). **(F)** Whole protein mass spectrometry of TEAD4 and TEAD4^C367S^ preincubated with 100 µM compound **1** (ACR-021) for 24 h at 4°C. TEAD4 peak was found at 25952 Da. Fragment adduct was detected at 26236 Da, an increase of 284 Da over protein alone matching the molecular weight of **1** (ACR-021). Fragment **1** (ACR-021) had additional peaks at Δ242 Da. TEAD4^C367S^ peak was found at 25936 Da and the compound did not form an adduct. **(G)** Whole protein mass spectrometry of TEAD4 and TEAD4^C367S^ preincubated with 100 µM compound **2** (ACR-362) for 24 h at 4°C. Fragment adducts were detected at Δ242 Da, matching the molecular weight of **2** (ACR-362). The fragment did not form an adduct with TEAD4^C367S^.

Subsequent studies focused on fragments **1** (ACR-021) and **2** (ACR-362) as they were the only fragments that inhibited TEAD binding to YAP through covalent bond formation at the central pocket cysteine. These two fragments were further tested in a concentration-dependent manner for inhibition of FAM-YAP_60–99_ binding to TEAD4 (**Fig. 1C**) and TEAD4^C367S^ mutant (**Fig. 1D**). Following 24 h incubation at 4°C, fragments **1** (ACR-021) and **2** (ACR-362) inhibited FAM-YAP_60–99_ peptide binding to wild-type TEAD4 (**Fig. 1C**) but not TEAD4^C367S^ mutant **(Fig. 1D**). Fragment **1** (ACR-021) inhibited wildtype TEAD4 with an IC_50_ of 4.6 ± 0.6 µM with maximum inhibition of 62%, but the fragment had no effect on TEAD4^C367S^. Fragment **2** (ACR-362) inhibited TEAD4 with a higher IC_50_ of 33.0 ± 6.0 µM. Fragments **1** (ACR-021) and **2** (ACR-362) were irreversible inhibitors of TEAD4 binding to YAP (FAM-YAP_60–99_) as established by a dialysis study (**Fig. 1E**).

Compound reaction to TEAD4 was analyzed by whole protein mass spectrometry (**Fig. 1F** and **G**). TEAD4 and TEAD4^C367S^ were incubated with 100 µM fragment for 24 h at 4°C. TEAD4 protein was found at 25952 Da and TEAD4^C367S^ mutant was found at 25936 Da. A TEAD4-**1** (ACR-021) adduct was found at 26236 Da with a difference of 284 Da, matching the molecular weight of **1** (ACR-021). Additional adducts were detected with adduct sizes of 242 mass units (**Fig. 1F**). Re-synthesis and testing of **1** (ACR-021) resulted in the same mass spectrometry pattern. Fragment

**2** (ACR-362) formed multiple adducts with TEAD4 with adduct sizes of 242 Da (**Fig. 1G**). At 100 µM, fragment **2** (ACR-362) formed three adducts with TEAD4. The compound did not react with TEAD4^C367S^ after 24 h at 4°C. Fragments **3**-**7**, **9** (ACR-195) and **10** (ACR-199) did not form detectable covalent adducts with TEAD4 or TEAD4^C367S^ (**Fig. S2C**). Fragment **8** (ACR-184) reacted to wildtype TEAD4 and formed two covalent adducts. Fragment **11** (ACR-300) formed a single adduct with wild-type TEAD4 but the adduct mass of 320 Da did not match the fragment expected molecular weight of 248.4 Da. Incubation with **12** (ACR-338) and **13** (ACR-345) led to formation of adducts of 107 Da (**Fig. S2C**). The 107 Da adduct matched the formation of adduct with dihydroquinone, which is a stabilizer molecule that was included in the library of acrylamide compounds by the manufacturer.

### Derivatives of 1 (ACR-021) Inhibit TEAD4

We selected **1** (ACR-021) for further exploration of the scaffold. Four derivatives, namely **14** (ACR-374) to **17** (ACR-375) (**Fig. 2A**), were prepared and evaluated for adduct formation and inhibition. The acrylamide warhead on the pyrrolidine ring of **1** (ACR-021) was rigidified as in **14** (ACR-374). The *para*-trifluoromethyl was moved to *meta*-[**15** (ACR-378)] and *ortho* positions [**16** (ACR-380)]. We also prepared **17** (ACR-375) with an octahydro-1*H*-pyrrolo[3,4-b]pyrazine ring. Derivatives of **1** (ACR-021) were tested for inhibition of FAM-YAP_60–99_ binding to TEAD4 or TEAD4^C367S^ (**Fig. 2B** and **C**). Compound **14** (ACR-374) inhibited wildtype TEAD4 with an IC_50_ of 2.9 ± 0.4 µM with maximum inhibition of 54% following 24 h incubation at 4°C (**Fig. 2B**). Moving the *para*-trifluoromethyl of **14** (ACR-374) to the *meta-*position as in **15** (ACR-378) and *ortho-*position as in **16** (ACR-380) led to reduced inhibition. Compound **17** (ACR-375) with the *para*-trifluoromethyl inhibited TEAD4 with a higher IC_50_ of 195 ± 20.4 µM. Both **14** (ACR-374) and **17** (ACR-375) inhibited wildtype TEAD4 but did not inhibit TEAD4^C367S^ (**Fig. 2C**). Parent fragment **1** (ACR-021) and its derivatives **14** (ACR-374) to **17** (ACR-375) did not inhibit TEAD1-3 binding to YAP (**Fig. 2D**-**F**).

**Figure 2.**
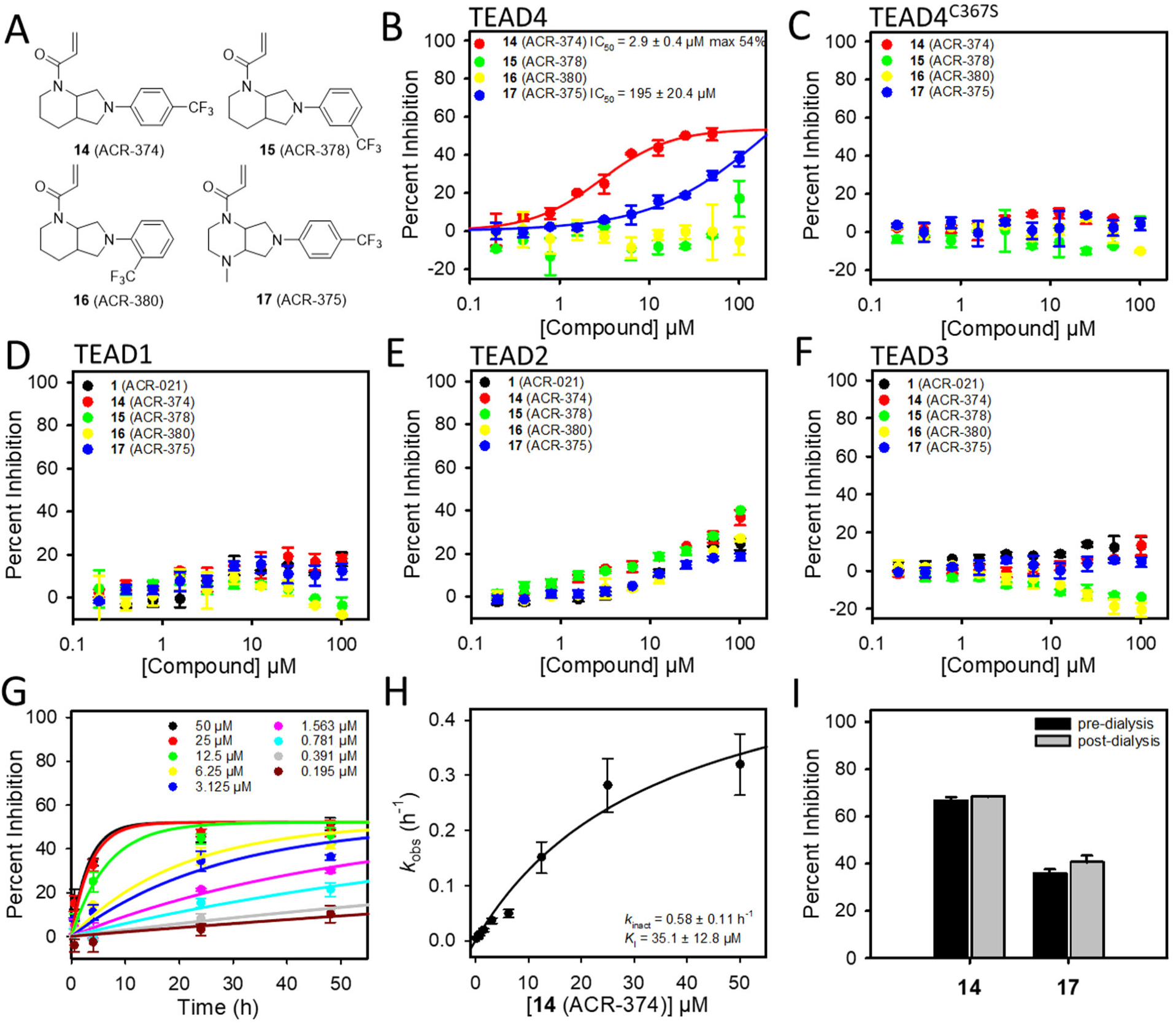
Inhibition of TEAD•YAP Binding by Derivatives of **1** (ACR-021). **(A)** Chemical structures of synthetic derivatives of **1** (ACR-021). **(B)** TEAD4 **(C)** TEAD4^C367S^ **(D)** TEAD1 **(E)** TEAD2 **(F)** TEAD3 were incubated with 0.2-100 µM derivatives for 24 h at 4°C and binding to fluorescently labeled YAP peptide (residues 60-99) was tested. Percent Inhibition was calculated relative to DMSO control and a sample without TEAD (mean ± s.d., n = 3). **(G)** Time-dependent inhibition of TEAD4 binding to YAP peptide by **14** (ACR-374). TEAD4 was incubated with 0.2-50 µM of **14** (ACR-374) and binding to YAP peptide was detected after 0.5, 6, 24 and 48 h incubation at 4°C. Inhibition versus time plots were fitted with exponential function to determine pseudo first-order rate constant *k*_obs_. **(H)** The *k*_obs_ were plotted against their respective concentrations of **14** (ACR-374) and fitted with a hyperbolic function 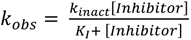 to determine *k*_inact_ and *K*_I_ values. **(I)** TEAD4 was incubated with 100 µM **14** (ACR-374) and **17** (ACR-375) for 24 h at 4°C. The samples were then dialyzed for 24 h at 4°C. Detection of binding to YAP peptide was performed before and after dialysis of the samples (n = 2).

A time- and concentration-dependent study determined the inactivation rate constant (*k*_inact_) and binding constant *K*_I_ for inhibition of the TEAD4•YAP protein-protein interaction. TEAD4 was incubated with varying concentrations of **14** (ACR-374) for 0.5, 6, 24 and 48 h at 4°C prior to binding studies using FAM-YAP_60–99_ peptide (**Fig. 2G**). The pseudo first-order rate constants (*k*_obs_) were determined by fitting an exponential function. The *k*_obs_ were plotted against their respective concentrations of **14** (ACR-374) and fitted with a hyperbolic function 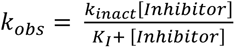 to determine the *k*_inact_ and *K*_I_ (**Fig. 2H**). The *k*_inact_ of inhibition was 0.45 ± 0.11 h^-1^ and the *K*_I_ was 61.3 ± 12.8 µM, leading to a second-order rate constant *k*_inact_/*K*_I_ of 2.04 ± 0.43 M^-1^s^-1^. Based on the *k*_inact_, the 𝑡^oo^ is 1.54 ± 0.14 h. Dialysis experiments were carried out and showed that, like **1** (ACR-021), **14** (ACR-374) and **17** (ACR-375) were also irreversible inhibitors of TEAD4 (**Fig. 2I**).

### Reaction and Inhibition Profile of 1 (ACR-021) and Derivatives Across All Four TEADs

Fragment **1** (ACR-021) and its derivatives **14** to **17** were tested for reaction to TEAD1-3 (**Fig. S3**), TEAD4, and TEAD4^C367S^ mutant (**Fig. 3A**). TEAD4 and TEAD4^C367S^ adducts were found at 25952 Da and 25936 Da, respectively. TEAD1, TEAD2 and TEAD3 were found at 27678 Da, 28431 Da and 27681 Da, respectively. For TEAD1-3 we observed two peaks for the apo proteins. The second peak was larger by 178 Da, which we established as gluconoylation at the N-terminal His-tag (41). Compounds **14** (ACR-374), **15** (ACR-378) and **16** (ACR-380) formed adducts as evidenced by a peak that is 324 Da larger than the protein alone. Incubation of protein with compound **17** (ACR-375) led to a peak that is 339 Da larger than the protein alone, matching the compound molecular weight. After incubating 100 µM compound for 24 h, the *para*-trifluoromethyl **14** (ACR-374) and *meta*-trifluoromethyl **15** (ACR-378) formed robust adducts with all TEADs, while the *ortho*-trifluoromethyl **16** (ACR-380) formed smaller or no adducts (**Table 1**).

**Figure 3.**
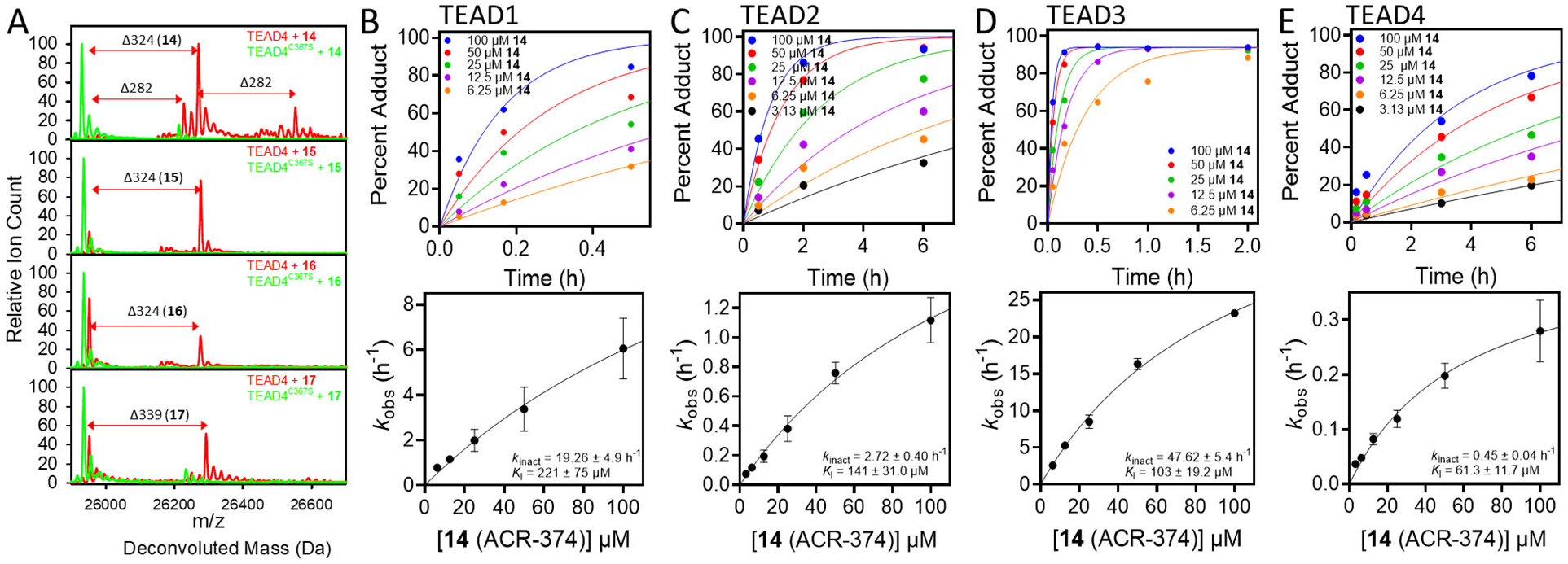
Reaction of **1** (ACR-021) Derivatives with TEADs. **(A)** TEAD4 and TEAD4^C367S^ mutant were incubated with 100 µM of **14**-**17** for 24 h at 4°C. TEAD4 protein peak was found at 25952 Da and adducts of **14**-**16** were found at 26276 Da with an adduct size of 324 Da. Compound **17** (ACR-375) formed an adduct at 26291 Da with an adduct size of Δ339 Da. Compound **14** (ACR-374) formed additional small peaks at Δ282 Da. TEAD4^C367S^ was found at 25936 Da and no adduct formation was detected. **(B)** TEAD1 was incubated with 6.25-100 µM of **14** (ACR-374) for 0.05-0.5 h at 4°C, and the reaction was quenched with 0.1 M formic acid. Whole-protein mass spectrometry was used to quantify the percent of TEAD1 covalently bound to **14** (ACR-374). Rate constants, *k*_obs_, for each concentration of **14** (ACR-374) were determined by fitting an exponential function. The *k*_obs_ were plotted against their respective concentrations, and a hyperbolic function 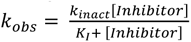 was fitted to calculate *k_inact_* and *K_I_*. **(C)** TEAD2 was incubated with 3.13-100 µM of **14** (ACR-374) for 0.5-6 h at 4°C, and the reaction was quenched with 0.1 M formic acid. Whole-protein mass spectrometry was used to quantify the percent of TEAD2 covalently bound to **14** (ACR-374). Rate constants, *k*_obs_, for each concentration of **14** (ACR-374) were determined by fitting an exponential function. The *k*_obs_ were plotted against their respective concentrations, and the hyperbolic function was fitted to calculate *k*_inact_ and *K_I_*. **(D)** TEAD3 was incubated with 6.25-100 µM of **14** (ACR-374) for 0.05-2 h at 4°C, and the reaction was quenched with 0.1 M formic acid. Whole-protein mass spectrometry was used to quantify the percent of TEAD3 covalently bound to **14** (ACR-374). Rate constants, *k*_obs_, for each concentration of **14** (ACR-374) were determined by fitting an exponential function. The *k*_obs_ were plotted against their respective concentrations, and the hyperbolic function was fitted to calculate *k*_inact_ and *K_I_*. **(E)** TEAD4 was incubated with 3.13-100 µM of **14** (ACR-374) for 0.05-6 h at 4°C, and the reaction was quenched with 0.1 M formic acid. Whole-protein mass spectrometry was used to quantify the percent of TEAD4 covalently bound to **14** (ACR-374). Rate constants, *k*_obs_, for each concentration of **14** (ACR-374) were determined by fitting an exponential function. The *k*_obs_ were plotted against their respective concentrations, and the hyperbolic function was fitted to calculate *k*_inact_ and *K_I_*.

**Table 1.**
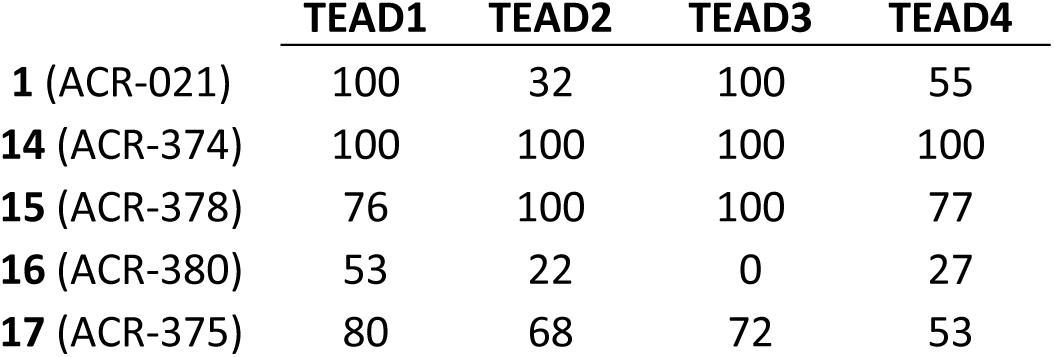
Percent Adduct Formation of 100 µM fragment at 24 h 4 °C.

Whole protein mass spectrometry was carried out in a time- and concentration-dependent manner to get insight into the reaction rates and binding constants of parental fragment and derivatives for each of the four mammalian TEADs. TEAD1-4 were incubated with varying concentrations of **14** (ACR-374) and aliquots were quenched at specific time points. The covalent complex formation was quantified by mass spectrometry and plotted against time (**Fig. 3B-E**). The pseudo first-order rate constants (*k*_obs_) were determined by fitting an exponential function for each concentration of **14** (ACR-374). The *k*_obs_ were plotted against their respective compound concentrations and were fitted with a hyperbolic equation 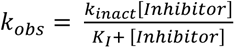 to determine *k*_inact_ and *K*_I._

The *k*_inact_ and *K*_I_ values for covalent bond formation with all four TEADs were determined for **1** (ACR-021), **2** (ACR-362) and **15** to **17** (**Fig. S4**) and the results are listed in **Table 2**. The second-order rate constant *k*_inact_/*K*_I_ is a good measure of overall covalent inhibitor potency, but 𝑡^oo^ and *K*_I_ provide deeper insight into the mechanism of inhibition. Fragment **1** (ACR-021) engaged TEAD3 with a *k*_inact_/*K*_I_ of 41.4 ± 3.1 M^-1^ s^-1^, while it was a 7.1 M^-1^ s^-1^ towards TEAD1. The reaction of **1** (ACR-021) with TEAD3 was fast with a 𝑡^oo^ in the minutes range compared to 9 hours for TEAD2. The *k*_inact_/*K*_I_ of **1** (ACR-021) for covalent bond formation with TEAD2 and TEAD4 was less than 1 M^-1^ s^-1^. Fragment **2** (ACR-362) had a low *k*_inact_/*K*_I_ for all four TEADs with 𝑡^oo^ values generally in the double-digit hours range. Compound **14** (ACR-374) reacted well with TEAD1 and TEAD3 with *k*_inact_/*K*_I_ values of 24.2 ± 10.3 M^-1^ s^-1^ and 128 ± 28 M^-1^ s^-1^, respectively. This is attributed mainly to substantial improvements in *k*_inact_, which correspond to 𝑡^oo^ values in the single-digit minute range or less. Like **1** (ACR-021), **14** (ACR-374) covalent bond formation was substantially more efficient with TEAD1 and TEAD3, compared with TEAD2 and TEAD4. A substantial improvement in 𝑡^oo^ from nearly 9 hours in the parent fragment to 30 minutes in **14** (ACR-374) was observed. Moving the *para*-trifluoromethyl moiety of **14** (ACR-374) to *meta* as in **15** (ACR-378) resulted in less favorable *k*_inact_/*K*_I_ mainly in the single-digit M^-1^ s^-1^ range. The *ortho*-trifluoromethyl of **16** (ACR-380) resulted in less than 1 M^-1^ s^-1^ *k*_inact_/*K*_I_ toward all TEADs. Similar results were seen for **17** (ACR-375), which had less than 1 M^-1^ s^-1^ *k*_inact_/*K*_I_ toward all TEADs.

**Table 2.**
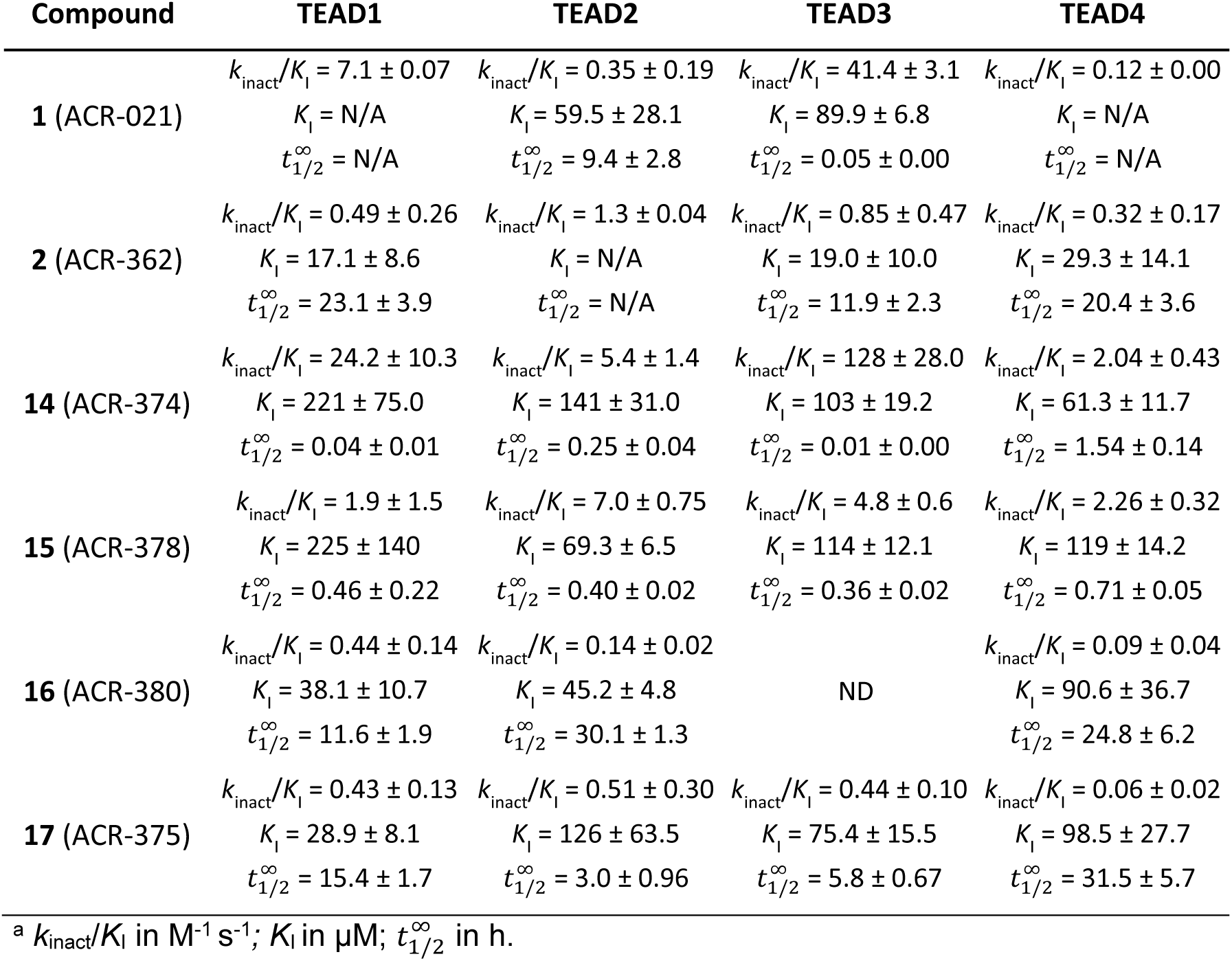
Reaction Constants of Acrylamides Toward TEADs ^a^.

| Compound | TEAD1 | TEAD2 | TEAD3 | TEAD4 |
| --- | --- | --- | --- | --- |
| **1** (ACR-021) | $k_{\text{inact}}/K_i = 7.1 \pm 0.07$ | $k_{\text{inact}}/K_i = 0.35 \pm 0.19$ | $k_{\text{inact}}/K_i = 41.4 \pm 3.1$ | $k_{\text{inact}}/K_i = 0.12 \pm 0.00$ |
| | $K_i = \text{N/A}$ | $K_i = 59.5 \pm 28.1$ | $K_i = 89.9 \pm 6.8$ | $K_i = \text{N/A}$ |
| | $t_{1/2}^{\infty} = \text{N/A}$ | $t_{1/2}^{\infty} = 9.4 \pm 2.8$ | $t_{1/2}^{\infty} = 0.05 \pm 0.00$ | $t_{1/2}^{\infty} = \text{N/A}$ |
| **2** (ACR-362) | $k_{\text{inact}}/K_i = 0.49 \pm 0.26$ | $k_{\text{inact}}/K_i = 1.3 \pm 0.04$ | $k_{\text{inact}}/K_i = 0.85 \pm 0.47$ | $k_{\text{inact}}/K_i = 0.32 \pm 0.17$ |
| | $K_i = 17.1 \pm 8.6$ | $K_i = \text{N/A}$ | $K_i = 19.0 \pm 10.0$ | $K_i = 29.3 \pm 14.1$ |
| | $t_{1/2}^{\infty} = 23.1 \pm 3.9$ | $t_{1/2}^{\infty} = \text{N/A}$ | $t_{1/2}^{\infty} = 11.9 \pm 2.3$ | $t_{1/2}^{\infty} = 20.4 \pm 3.6$ |
| **14** (ACR-374) | $k_{\text{inact}}/K_i = 24.2 \pm 10.3$ | $k_{\text{inact}}/K_i = 5.4 \pm 1.4$ | $k_{\text{inact}}/K_i = 128 \pm 28.0$ | $k_{\text{inact}}/K_i = 2.04 \pm 0.43$ |
| | $K_i = 221 \pm 75.0$ | $K_i = 141 \pm 31.0$ | $K_i = 103 \pm 19.2$ | $K_i = 61.3 \pm 11.7$ |
| | $t_{1/2}^{\infty} = 0.04 \pm 0.01$ | $t_{1/2}^{\infty} = 0.25 \pm 0.04$ | $t_{1/2}^{\infty} = 0.01 \pm 0.00$ | $t_{1/2}^{\infty} = 1.54 \pm 0.14$ |
| **15** (ACR-378) | $k_{\text{inact}}/K_i = 1.9 \pm 1.5$ | $k_{\text{inact}}/K_i = 7.0 \pm 0.75$ | $k_{\text{inact}}/K_i = 4.8 \pm 0.6$ | $k_{\text{inact}}/K_i = 2.26 \pm 0.32$ |
| | $K_i = 225 \pm 140$ | $K_i = 69.3 \pm 6.5$ | $K_i = 114 \pm 12.1$ | $K_i = 119 \pm 14.2$ |
| | $t_{1/2}^{\infty} = 0.46 \pm 0.22$ | $t_{1/2}^{\infty} = 0.40 \pm 0.02$ | $t_{1/2}^{\infty} = 0.36 \pm 0.02$ | $t_{1/2}^{\infty} = 0.71 \pm 0.05$ |
| **16** (ACR-380) | $k_{\text{inact}}/K_i = 0.44 \pm 0.14$ | $k_{\text{inact}}/K_i = 0.14 \pm 0.02$ | | $k_{\text{inact}}/K_i = 0.09 \pm 0.04$ |
| | $K_i = 38.1 \pm 10.7$ | $K_i = 45.2 \pm 4.8$ | ND | $K_i = 90.6 \pm 36.7$ |
| | $t_{1/2}^{\infty} = 11.6 \pm 1.9$ | $t_{1/2}^{\infty} = 30.1 \pm 1.3$ | | $t_{1/2}^{\infty} = 24.8 \pm 6.2$ |
| **17** (ACR-375) | $k_{\text{inact}}/K_i = 0.43 \pm 0.13$ | $k_{\text{inact}}/K_i = 0.51 \pm 0.30$ | $k_{\text{inact}}/K_i = 0.44 \pm 0.10$ | $k_{\text{inact}}/K_i = 0.06 \pm 0.02$ |
| | $K_i = 28.9 \pm 8.1$ | $K_i = 126 \pm 63.5$ | $K_i = 75.4 \pm 15.5$ | $K_i = 98.5 \pm 27.7$ |
| | $t_{1/2}^{\infty} = 15.4 \pm 1.7$ | $t_{1/2}^{\infty} = 3.0 \pm 0.96$ | $t_{1/2}^{\infty} = 5.8 \pm 0.67$ | $t_{1/2}^{\infty} = 31.5 \pm 5.7$ |

<sup>a</sup> $k_{\text{inact}}/K_i$ in $\text{M}^{-1} \text{s}^{-1}$ ; $K_i$ in $\mu\text{M}$ ; $t_{1/2}^{\infty}$ in h.

### Crystal Structures of TEAD2 and TEAD3 in Complex with Acrylamide Fragments

Covalent complex crystal structures were obtained by soaking apo TEAD2 crystals with fragments **14**-**16**. Two monomers of TEAD2 were found in the asymmetric unit, monomers A and B. TEAD2-**14** (ACR-374) complex structure was solved at 2.60 Å resolution (**Table S2**). Each monomer within the asymmetric unit contained a **14** (ACR-374) in a covalent complex with TEAD. In monomer A, **14** (ACR-374) is in a covalent complex with TEAD2 Cys-380 (**Fig. 4A**). The density of the compound was strong around the covalent bond and the trifluoromethylphenyl moiety, but it was weaker around the octahydro-1*H*-pyrrolo[3,4-b]pyridine (**Fig. 4B**). The strong density around the trifluoromethylphenyl indicated the moiety to be well positioned in the sub-pocket surrounded by hydrophobic Ala-235, Ile-408 and Val-252 as well as aromatic Phe-428, Phe-233, Phe-406 and Tyr-426 (**Fig. 4A**). The octahydro-1*H*-pyrrolo[3,4-b]pyridine is located next to Ser-345 and Val-329.

**Figure 4.**
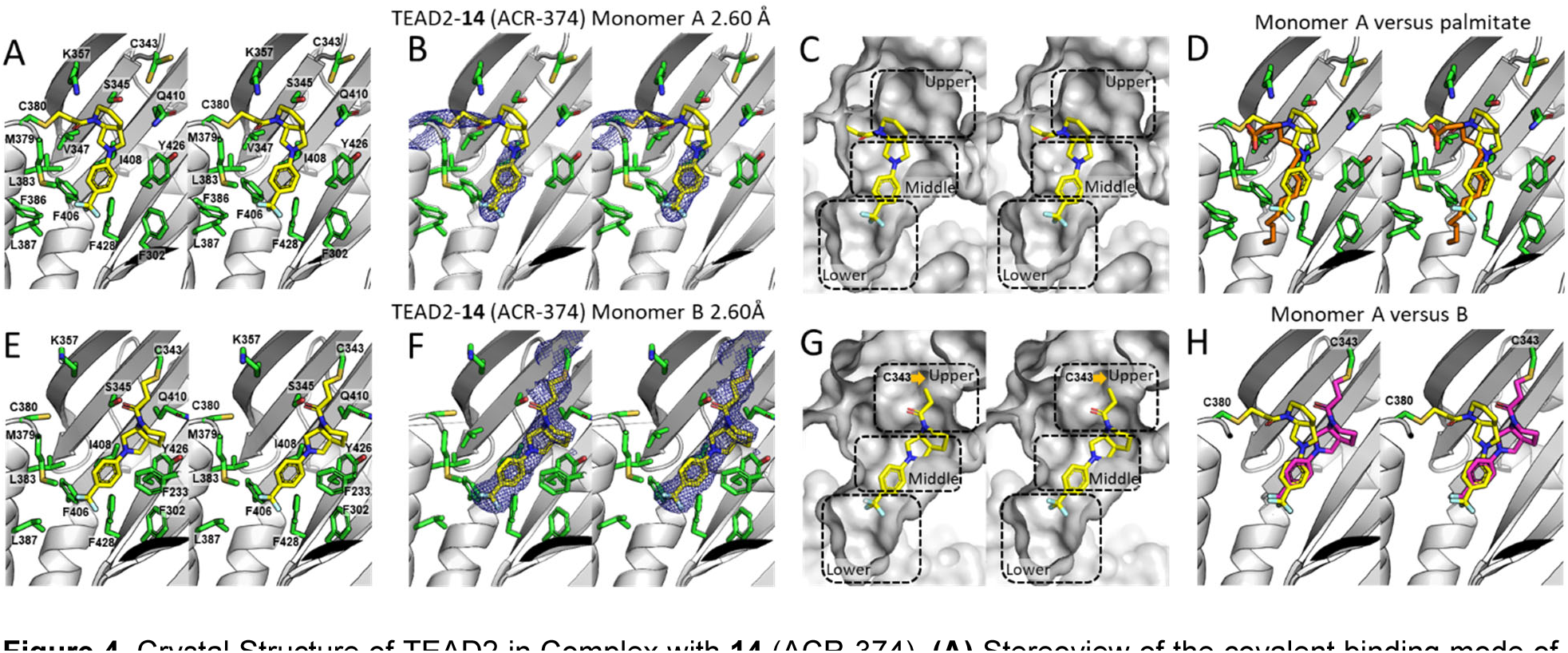
Crystal Structure of TEAD2 in Complex with **14** (ACR-374). **(A)** Stereoview of the covalent binding mode of **14** (ACR-374) [capped-sticks representation with carbon, nitrogen, oxygen, and fluorine in yellow, blue, red, and cyan, respectively] in complex with TEAD2 in monomer A (gray ribbon representation). TEAD2 residues in the pocket are shown in a capped-sticks representation (carbon, nitrogen, oxygen, and sulfur in green, blue, red, and gold, respectively). **(B)** Stereoview of the composite omit map around TEAD2 Cys-380 and **14** (ACR-374) in monomer A. The map is shown in blue mesh at σ 1.0. **(C)** Stereoview of the covalent binding mode of **14** (ACR-374) [capped-sticks representation; carbon, nitrogen, oxygen, and fluorine in yellow, blue, red, and cyan, respectively] in complex with TEAD2 monomer A (gray Connolly surface representation). The sub-pockets within are highlighted with dashed lines. **(D)** Stereoview of the covalent binding mode of **14** (ACR-374) [capped-sticks with carbon, nitrogen, oxygen, and fluorine in yellow, blue, red, and cyan, respectively] in TEAD2 monomer A overlapped with TEAD2-bound palmitate (capped-sticks with carbon and oxygen in orange and red, respectively) [PDB: 5HGU]. **(E)** Stereoview of the covalent binding mode of **14** (ACR-374) [capped-sticks representation with carbon, nitrogen, oxygen, and fluorine in yellow, blue, red, and cyan, respectively] in complex with TEAD2 in monomer B (gray ribbon representation). TEAD2 residues in the pocket are shown in a capped-sticks representation (carbon, nitrogen, oxygen, and sulfur in green, blue, red, and gold, respectively). **(F)** Stereoview of the composite omit map around TEAD2 Cys-343 and **14** (ACR-374) in monomer B. The map is shown in blue mesh at σ 1.0. **(G)** Stereoview of the covalent binding mode of **14** (ACR-374) [capped-sticks representation; carbon, nitrogen, oxygen, and fluorine in yellow, blue, red, and cyan, respectively] in complex with TEAD2 monomer B (gray Connolly surface representation). The sub-pockets within are highlighted with dashed lines. **(H)** Stereoview of the covalent binding mode of **14** (ACR-374) in TEAD2 monomer A (capped-sticks with carbon, nitrogen, oxygen, and fluorine in yellow, blue, red, and cyan, respectively) overlapped with the binding mode of **14** (ACR-374) in TEAD2 monomer B (capped-sticks with carbon, nitrogen, oxygen, and fluorine in magenta, blue, red, and cyan, respectively).

The palmitate pocket in TEADs can be divided into three sub-pockets as shown in **Fig. 4C**. The native palmitate occupies the “middle” and “lower” sub-pockets and does not engage the “upper” sub-pocket. In TEAD2 monomer A, **14** (ACR-374) engages the middle sub-pocket with the trifluoromethyl moiety reaching into the lower sub-pocket. The piperidine ring of the octahydro-1*H*-pyrrolo[3,4-b]pyridine moiety engages the upper sub-pocket. Structural superimposition of TEAD2 in complex with palmitate (PDB: 5HGU) and TEAD2 in a covalent complex with **14** (ACR-374) shows that palmitate engages the lower sub-pocket fully with additional four carbon atoms beyond the structure of **14** (ACR-374) [**Fig. 4D**].

Unexpectedly, in monomer B, **14** (ACR-374) forms a covalent bond with Cys-343 of TEAD2 (**Fig. 4E**). The electron density for this structure is well defined in monomer B (**Fig. 4F**). Despite the reaction at different cysteines, the location of the trifluoromethylphenyl moiety of the compound in monomer B is in the same position as that of monomer A. The octahydro-1*H*-pyrrolo[3,4-b]pyridine moiety is closer to Tyr-436, Gln-410, Ser-345 and Phe-233. Gln-410 adopts a different conformation. As with the binding mode observed in monomer A, most of the compound is in the middle sub-pocket (**Fig. 4G**). The trifluoromethyl points into the lower sub-pocket and the warhead, which forms a covalent bond with Cys-343, is located in the upper sub-pocket. Superimposition of **14** (ACR-374) in the two monomers (**Fig. 4H**) shows the trifluoromethylphenyl moiety is positioned at the same place, while the rest of the molecule moves between pointing toward Cys-380 and Cys-343.

TEAD2-**15** (ACR-378) complex structure was solved at 2.72 Å resolution (**Table S2**). Both monomers contained **15** (ACR-378) in a covalent bond with Cys-380 (**Fig. 5A** and **E**). The electron density of **15** (ACR-378) in monomer A covered the whole molecule (**Fig. 5B**) while that of monomer B was weaker around the octahydro-1*H*-pyrrolo[3,4-b]pyridine (**Fig. 5F**). In both monomers, the density around the trifluoromethylphenyl was strong. As with the binding mode of **14** (ACR-374) in monomer A, **15** (ACR-378) in monomer A mostly occupies the middle sub-pocket with the trifluoromethyl pointing into the lower pocket, and the piperidine ring pointing towards the upper pocket (**Fig. 5C**). Superimposition of the binding mode of **14** (ACR-374) in monomer A onto that of **15** (ACR-378) shows the fragments in a similar position. The *meta*-trifluoromethyl of **15** (ACR-378) does not engage as deeply into the lower sub-pocket. In monomer B, the meta-trifluoromethyl moiety allows for the compound to point toward the TEAD2 Cys-380 residue, avoiding the linear binding mode of **14** (ACR-374) observed in monomer B. The overall binding modes of **15** (ACR-378) in monomers A and B are similar (**Fig. 5G** and **H**).

**Figure 5.**
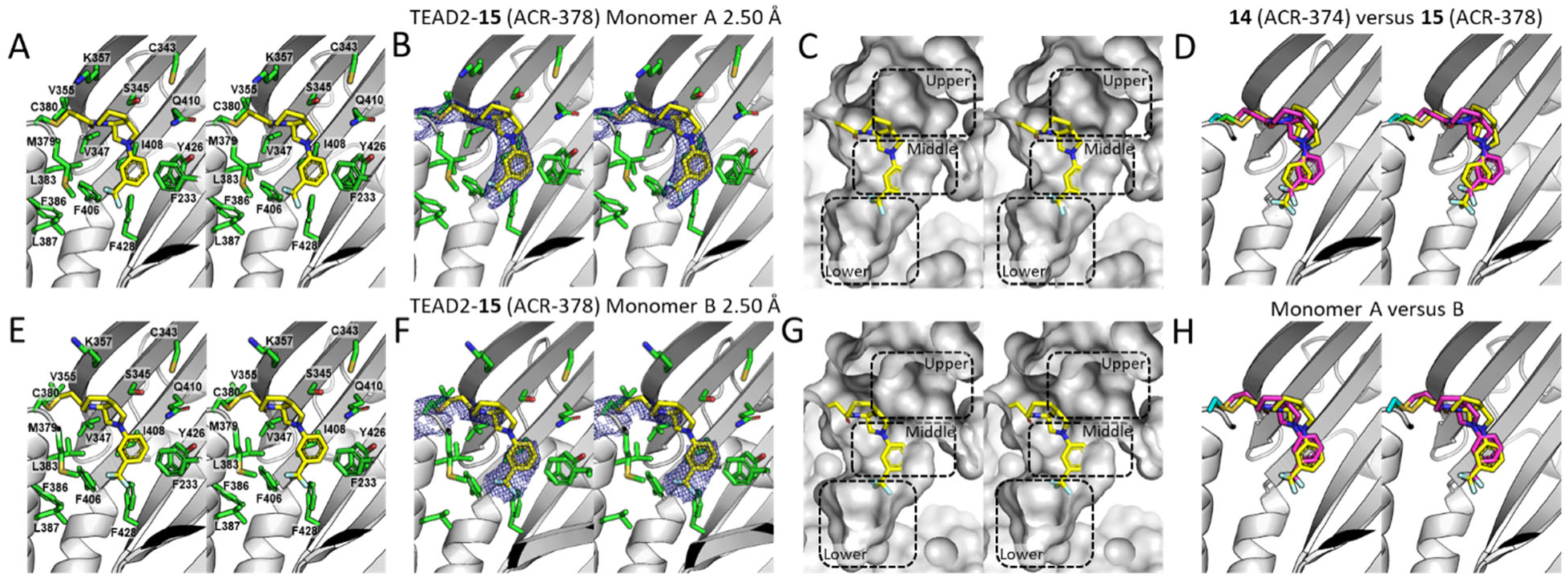
Crystal Structure of TEAD2 in Complex with **15** (ACR-378). **(A)** Stereoview of the covalent binding mode of **15** (ACR-378) [capped-sticks representation with carbon, nitrogen, oxygen, and fluorine in yellow, blue, red, and cyan, respectively] in complex with TEAD2 in monomer A (gray ribbon representation). TEAD2 residues in the pocket are shown in a capped-sticks representation (carbon, nitrogen, oxygen, and sulfur in green, blue, red, and gold, respectively). **(B)** Stereoview of the composite omit map around TEAD2 Cys-380 and **15** (ACR-378) in monomer A. The map is shown in blue mesh at σ 1.0. **(C)** Stereoview of the covalent binding mode of **15** (ACR-378) [capped-sticks representation; carbon, nitrogen, oxygen, and fluorine in yellow, blue, red, and cyan, respectively] in complex with TEAD2 monomer A (gray Connolly surface representation). The sub-pockets within are highlighted with dashed lines. **(D)** Stereoview of the covalent binding mode of **15** (ACR-378) [capped-sticks with carbon, nitrogen, oxygen, and fluorine in magenta, blue, red, and cyan, respectively] in TEAD2 monomer A overlapped with the covalent binding mode of **14** (ACR-374) [capped-sticks with carbon, nitrogen, oxygen, and fluorine in yellow, blue, red, and cyan, respectively]. **(E)** Stereoview of the covalent binding mode of **15** (ACR-378) [capped-sticks representation with carbon, nitrogen, oxygen, and fluorine in yellow, blue, red, and cyan, respectively] in complex with TEAD2 in monomer B (gray ribbon representation). TEAD2 residues in the pocket are shown in a capped-sticks representation (carbon, nitrogen, oxygen, and sulfur in green, blue, red, and gold, respectively). **(F)** Stereoview of the composite omit map around TEAD2 Cys-380 and **15** (ACR-378) in monomer B. The map is shown in blue mesh at σ 1.0. **(G)** Stereoview of the covalent binding mode of **15** (ACR-378) [capped-sticks representation; carbon, nitrogen, oxygen, and fluorine in yellow, blue, red, and cyan, respectively] in complex with TEAD2 monomer B (gray Connolly surface representation). The sub-pockets within are highlighted with dashed lines. **(H)** Stereoview of the covalent binding mode of **15** (ACR-378) in TEAD2 monomer A (capped-sticks with carbon, nitrogen, oxygen, and fluorine in yellow, blue, red, and cyan, respectively) overlapped with the binding mode of **15** (ACR-378) in TEAD2 monomer B (capped-sticks with carbon, nitrogen, oxygen, and fluorine in magenta, blue, red, and cyan, respectively).

The structure of TEAD2-**16** (ACR-380) complex was solved at 2.23 Å resolution (**Table S2**). The compound formed a covalent bond with Cys-380 in both monomer A and B (**Fig. 6A** and **D**). Density for the compound was found within both monomers (**Fig. 6B** and **E**). The binding mode of **16** (ACR-380) in both monomers reveals that the compound occupies mainly the middle pocket (**Fig. 6C** and **F**). Interestingly, the *ortho*-trifluoromethyl does not engage the lower pocket at all, compared to **14** (ACR-374) and **15** (ACR-378).

**Figure 6.**
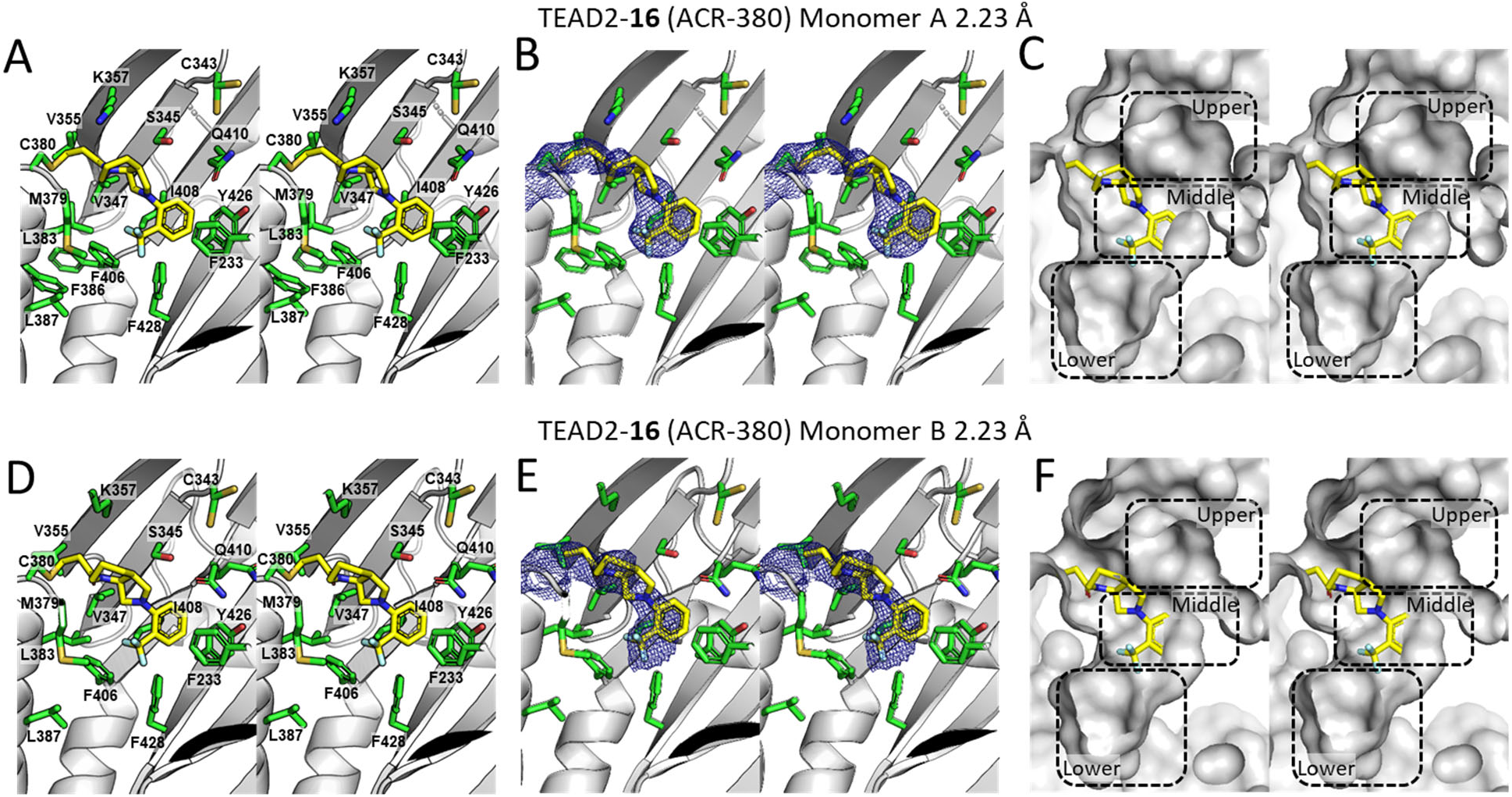
Crystal Structure of TEAD2 in Complex with **16** (ACR-380). **(A)** Stereoview of the covalent binding mode of **16** (ACR-380) [capped-sticks representation with carbon, nitrogen, oxygen, and fluorine in yellow, blue, red, and cyan, respectively] in complex with TEAD2 in monomer A (gray ribbon representation). TEAD2 residues in the pocket are shown in a capped-sticks representation (carbon, nitrogen, oxygen, and sulfur in green, blue, red, and gold, respectively). **(B)** Stereoview of the composite omit map around TEAD2 Cys-380 and **16** (ACR-380) in monomer A. The map is shown in blue mesh at σ 1.0. **(C)** Stereoview of the covalent binding mode of **16** (ACR-380) [capped-sticks representation; carbon, nitrogen, oxygen, and fluorine in yellow, blue, red, and cyan, respectively] in complex with TEAD2 monomer A (gray Connolly surface representation). The sub-pockets within are highlighted with dashed lines. **(D)** Stereoview of the covalent binding mode of **16** (ACR-380) [capped-sticks representation with carbon, nitrogen, oxygen, and fluorine in yellow, blue, red, and cyan, respectively] in complex with TEAD2 in monomer B (gray ribbon representation). TEAD2 residues in the pocket are shown in a capped-sticks representation (carbon, nitrogen, oxygen, and sulfur in green, blue, red, and gold, respectively). **(E)** Stereoview of the composite omit map around TEAD2 Cys-380 and **16** (ACR-380) in monomer B. The map is shown in blue mesh at σ 1.0. **(F)** Stereoview of the covalent binding mode of **16** (ACR-380) [capped-sticks representation; carbon, nitrogen, oxygen, and fluorine in yellow, blue, red, and cyan, respectively] in complex with TEAD2 monomer B (gray Connolly surface representation). The sub-pockets within are highlighted with dashed lines.

TEAD3-**14** (ACR-374) complex structure was obtained by co-crystallization and the structure was solved at 2.98 Å resolution (**Table S2**). Four monomers of TEAD3 were found in the asymmetric unit and each monomer contained **14** (ACR-374) in a covalent complex in the palmitate pocket (**Fig. 7A**). The electron density for the compound was strong in all four monomers with the best density found in monomer B (**Fig. 7B** and **Fig. S5**). Since the binding mode of **14** (ACR-374) in all four monomers was identical (**Fig. 7C**), structural analysis is performed on monomer B. The binding mode of **14** (ACR-374) in TEAD3 mimics the binding mode of palmitate (**Fig. 7D**). Remarkably, the trifluoromethylphenyl moiety fully occupies the lower pocket surrounded by residues Phe-247, Ile-375, Leu-303, Phe-416 and Phe-394 (**Fig. 7A** and **E**). The octahydro-1*H*-pyrrolo[3,4-b]pyridine moiety is found in the middle pocket, surrounded by Met-367, Met-371, Ile-396, Phe-414 and Tyr-230 (**Fig. 7A** and **E**). No part of the compound engages the upper pocket (**Fig. 7E**). The superimposition of the binding mode of **14** (ACR-374) from TEAD2 monomer A onto that of TEAD3 highlights the large difference of the binding modes (**Fig. 7F**). The residues in the middle and lower sub-pockets of TEAD2 and TEAD3 are similar with two differences. The TEAD2 Leu-383 equivalent in TEAD3 is Met-371, while TEAD2 Phe-233 is equivalent to TEAD3 Tyr-230. The distance between TEAD3 Met-371 and **14** (ACR-374) is 4.0 Å. Measuring the distance from TEAD2 Leu-383 to **14** (ACR-374) in TEAD3 shows 2.0 Å, indicating a clash would occur in TEAD2. The closest distance between TEAD2 Phe-233 to the **14** (ACR-374) bound to TEAD2 is 3.5 Å. The closest distance between TEAD3 Tyr-230 hydroxyl to the **14** (ACR-374) bound to TEAD2 is 2.0 Å, also indicating that a clash would occur. The subtle difference in the nature of the binding site residues of TEAD2 and TEAD3 led to drastically different binding modes of the same compound. The differences in binding modes resulted in **14** (ACR-374) having a *k*_inact_/*K*_I_ of 5.4 M^-1^ s^-1^ to TEAD2 while having a *k*_inact_/*K*_I_ of 128 M^-1^ s^-1^ to TEAD3 (**Table 2**).

**Figure 7.**
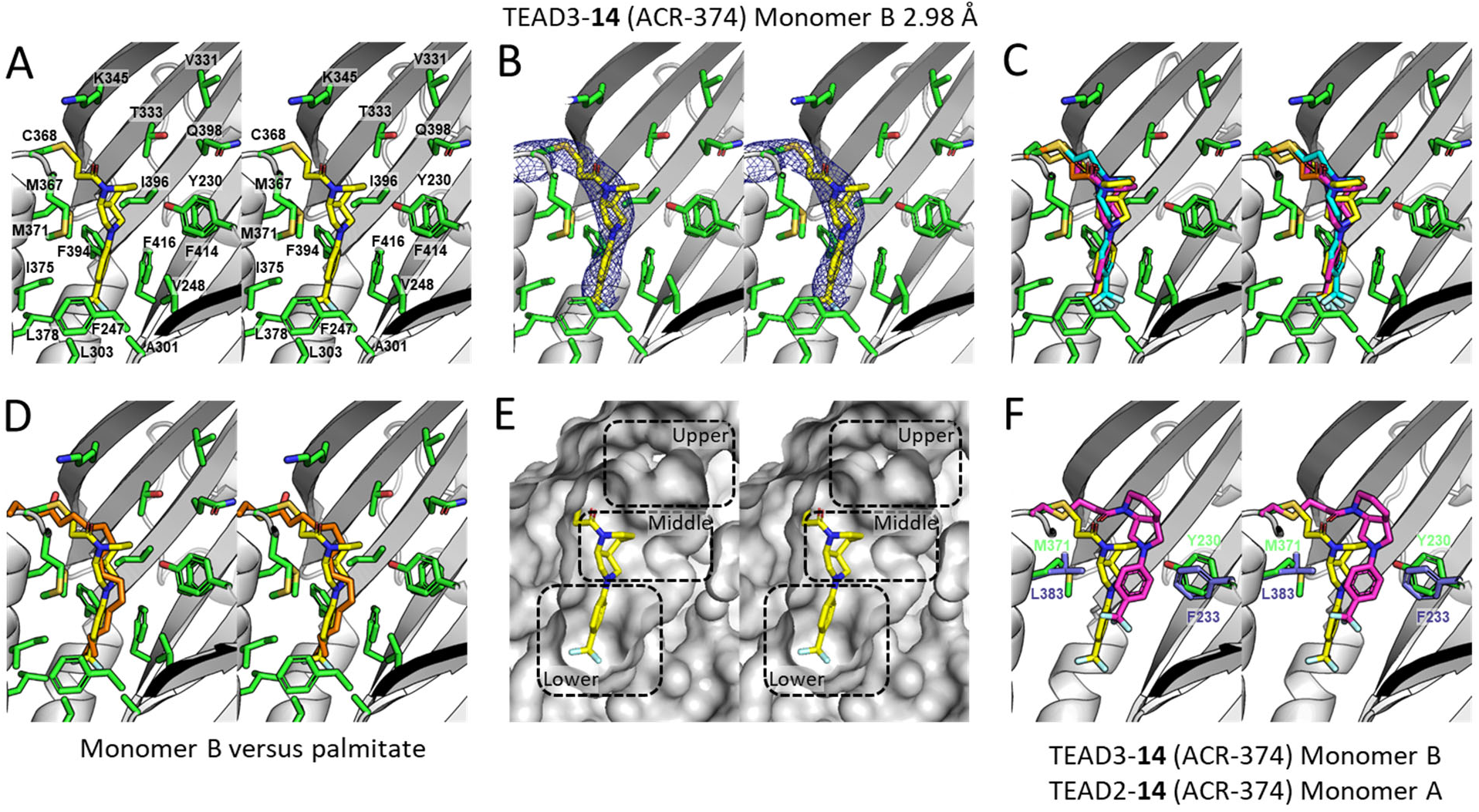
**Crystal Structure of TEAD3 in Complex with 14 (ACR-374)**. **(A)** Stereoview of the covalent binding mode of **14** (ACR-374) [capped-sticks representation with carbon, nitrogen, oxygen, and fluorine in yellow, blue, red, and cyan, respectively] in complex with TEAD3 in monomer B (gray ribbon representation). TEAD3 residues in the pocket are shown in a capped-sticks representation (carbon, nitrogen, oxygen, and sulfur in green, blue, red, and gold, respectively). **(B)** Stereoview of the composite omit map around TEAD3 Cys-368 and **14** (ACR-374) in monomer B. The map is shown in blue mesh at σ 1.0. **(C)** Stereoview of the covalently bound **14** (ACR-374) in TEAD3 monomers A-D. **(D)** Stereoview of the covalent binding mode **14** (ACR-374) [capped-sticks with carbon, nitrogen, oxygen, and fluorine in yellow, blue, red, and cyan, respectively] in TEAD3 monomer B overlapped with TEAD3-bound palmitate (capped-sticks with carbon and oxygen in orange and red, respectively) [PDB: 5EMW]. **(E)** Stereoview of the covalent binding mode of **14** (ACR-374) [capped-sticks representation; carbon, nitrogen, oxygen, and fluorine in yellow, blue, red, and cyan, respectively] in complex with TEAD3 monomer B (gray Connolly surface representation). The sub-pockets within are highlighted with dashed lines. **(F)** Stereoview of the covalent binding mode of **14** (ACR-374) [capped-sticks with carbon, nitrogen, oxygen, and fluorine in yellow, blue, red, and cyan, respectively] in TEAD3 monomer B overlapped with the covalent binding mode of **14** (ACR-374) [capped-sticks with carbon, nitrogen, oxygen, and fluorine in magenta, blue, red, and cyan, respectively] in TEAD2 monomer A. Side-chains of TEAD2 residues Phe-233 and Leu-383 are shown in capped-sticks representation (purple carbons). Side-chains of TEAD3 Tyr-230 and Met-371 are shown in capped-sticks representation (carbon, oxygen, and sulfur in green, red, and gold, respectively).

### Engagement of Lipid Pocket Amino Acids Alters Small-Molecule Inhibition Profile

The lipid pocket of TEADs is located outside the TEAD•YAP protein-protein interface. Inhibition of TEAD4 binding to YAP likely occurs through an allosteric mechanism. Allostery is a complex process that is highly system dependent. In some cases, allosteric effects may occur through specific channels whereby structural changes are transferred from one amino acid to the next starting at one site and ending at another site on the protein. Other mechanisms of allostery have been proposed that involve global conformational changes. Regardless of the mechanism, the allosteric effect is initiated at the distal binding pocket by the small molecule. Our hypothesis is that engagement of amino acids within the palmitate pocket by the small molecules leads to a series of conformational changes that alter the structure of TEAD4 resulting in lower affinity binding to YAP. To test this hypothesis, we mutated two amino acids within the lipid pocket of TEAD4, and we repeated the inhibition studies with these mutants using our fluorescence polarization assay for two compounds, namely **14** (ACR-374) and previously published TED-642 (**Fig. 8**). The two amino acids, namely TEAD4 Phe-393 and Phe-415, were selected for mutation using our crystal structure of **14** (ACR-374) bound to TEAD2 (**Fig. 8A**). These amino acids (Phe-406 and Phe-428 in TEAD2), come into direct contact with the compound. Interestingly, there were noticeable differences in the inhibition of TEAD4 mutant and TEAD4 wild-type binding to YAP for compounds **14** (ACR-374) [**Fig. 8B**] and TED-642 (**Fig. 8C**), respectively. The compounds inhibited TEAD4 mutant binding to YAP much better than wild-type TEAD4. The mutation of phenylalanine to alanine in each case resulted in a stronger allosteric effect. These results support our hypothesis that engagement of specific amino acids within the palmitate pocket is likely responsible for the distant structural changes that lead to inhibition of YAP binding to TEADs.

**Figure 8.**
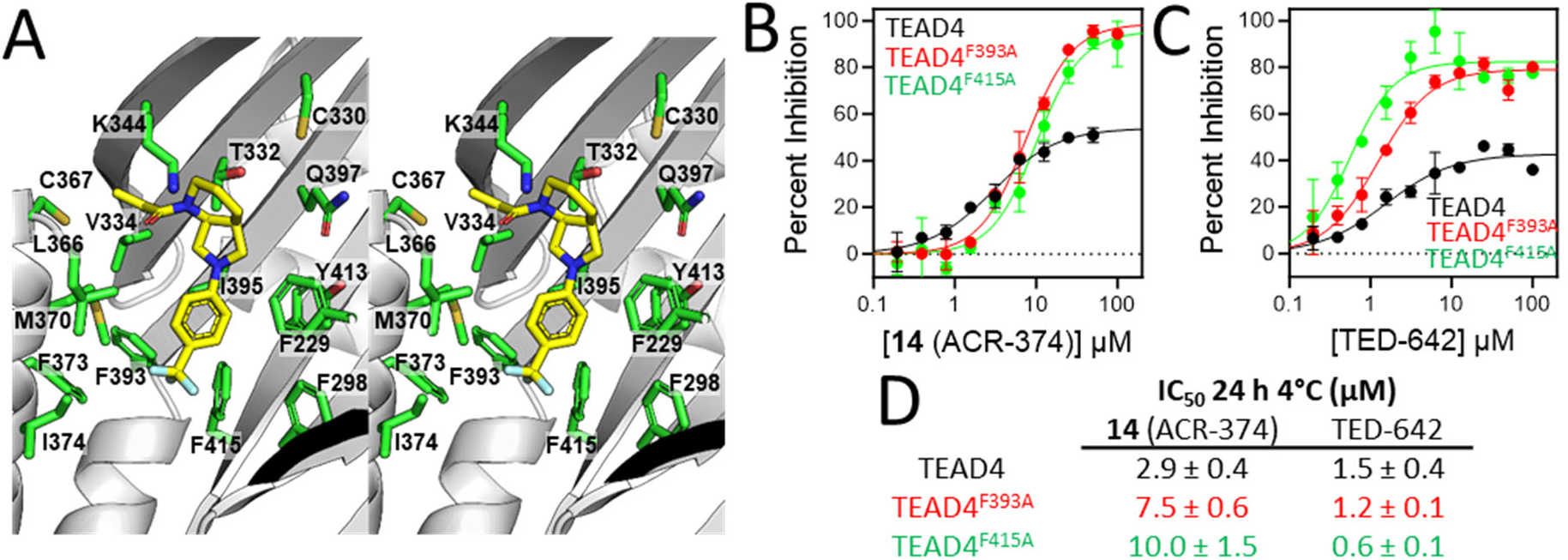
**Compound Binding to Amino Acids in the Palmitate Pocket Leads to Allosteric Inhibition**. **(A)** Stereoview of TEAD4 (gray ribbon representation) superimposed to TEAD2 crystal structure (not shown) bound to **14** (ACR-374) highlighting the interaction between the small molecule and pocket amino acids. Compound and amino acids are shown in capped-sticks representation with carbon in yellow for compound and green for amino acids, and oxygen, nitrogen, fluorine, sulfur in red, blue, cyan, and gold, respectively). **(B)** TEAD4^F393A^ and TEAD4^F415A^ mutants were preincubated with 0.2-100 µM **14** (ACR-374) for 24 h at 4°C before measuring binding to the fluorescently labeled YAP peptide. Percent inhibition was determined relative to DMSO control (n = 2). **(C)** TEAD4, TEAD4^F393A^ and TEAD4^F415A^ were preincubated with 0.2-100 µM TED-642 for 24 h at 4°C before measuring its binding to the fluorescently labeled YAP_60-99_ peptide (n = 2). **(D)** IC_50_ values at 24 h 4°C for **14** (ACR-374) and TED-642.

## DISCUSSION

We have previously shown that inhibition of TEAD•YAP is possible through binding in the central palmitate pocket of TEADs located outside the protein-protein interaction interface, and covalent reaction with a conserved cysteine in that pocket (17). Since then, numerous small molecules that bind to the TEAD binding pocket have been reported (**Figure S6**). To date, there is only one reported acrylamide small molecule that binds to the palmitate pocket and inhibits TEAD binding to YAP (32). We reported TED-642 that reacts at the cysteine in the palmitate pocket and inhibits TEAD binding to YAP (24). In other work, a series of vinyl sulfones, among them DC-TEADin02 and DC-TEAD3in03, inhibited palmitate binding, but they had no effect on TEAD binding to YAP (25, 26). Acrylamide K-975 was described as a covalent inhibitor of TEAD palmitoylation and *in vitro* transcriptional activity (27). In our hands, the compound did not inhibit TEAD1-4 binding to YAP in our fluorescence polarization assay (**Figure S7**). A series of highly reactive and unstable Kojic acids were also shown to react in the palmitate pocket and disrupt binding to YAP (28). Acrylamide MYF-01-37 reacted with TEAD palmitate cysteine in a mass spectrometry screen and inhibited TEAD in a cell-based luciferase assay (29, 30). The scaffold was further modified to MYF-03-176, which had nanomolar affinity in the palmitate pocket and showed significant inhibition of gene expression and proliferation in cell based assays (31). A set of non-covalent inhibitors of TEAD binding to YAP was turned into covalent inhibitor mCMY020, which displaced palmitate with nanomolar IC_50_ and inhibited cellular transcription and luciferase assays (32). Another acrylamide SWTX-143 was found to react with the palmitate cysteine, improve TEAD thermal stability, inhibit TEAD reporter assay and downregulate TEAD downstream gene transcription (33). In addition to covalent inhibitors, a number of interesting non-covalent inhibitor scaffolds were reported including one recent compound from Genentech that inhibited the TEAD•YAP protein-protein interaction (34–40).

Here, we resort to covalent fragment screening to identify acrylamide fragment starting points that could be used for the development of covalent TEAD•YAP inhibitors. We were interested not only in compounds that bind to the palmitate pocket and inhibit palmitate binding but also in compounds that inhibit TEAD binding to YAP. To that end, our screen was carried out using a fluorescence polarization assay that uses TEAD4 and a YAP peptide that spans the entire TEAD•YAP protein-protein interaction interface. Although several hits were identified, only fragments that inhibited TEAD4 binding to YAP through covalent bond formation at the central pocket cysteine were selected. Fragment **1** (ACR-021) was selected for further studies. Resynthesis confirmed its inhibition, and we designed a few derivatives such as **14** (ACR-374) to further establish the inhibition of the fragment and to establish limited structure-activity relationships and to gain further insight into the basis for inhibition of the protein-protein interaction.

Time- and concentration-dependent studies provide reaction rate *k*_inact_ and binding constant *K*_I_. From the reaction rate *k*_inact_, the reaction half-life t_1/2_ can be obtained providing insight into the reaction timescale. Time- and concentration-dependent studies using intact mass spectrometry provide information about binding and reactivity of compounds to TEADs. Time- and concentration-dependent studies using our fluorescence polarization assay probe the rates of inhibition of the protein-protein interaction. For **14** (ACR-374), the reaction and inhibition timescales were similar, with reaction half-life from mass spectrometry and fluorescence polarization of about an hour. This suggests that the conformational changes induced by covalent bond formation occur rapidly following the reaction. This means that the conformational changes are likely local structural changes of amino acid sidechains rather than global changes that may take longer to occur. Finally, the lack of complete inhibition of the protein-protein interaction suggests that the compound may not be inducing dramatic changes to the structure. The structural changes likely weaken the binding of TEAD to YAP, shifting the equilibrium towards less binding.

It is interesting that the compounds show much higher inhibition of TEAD4 binding to YAP than the other TEADs despite the high sequence conservation in the binding pocket. Some inhibition of TEAD2 binding to YAP is observed at higher concentrations near 100 µM by about 30% for **14** (ACR-374), but no inhibition for TEAD1 and TEAD3 binding to YAP up to 100 µM. The differences in binding modes of the compounds in TEAD2 and TEAD3 that we observe in our co-crystal structures potentially provide an explanation. In TEAD2, the compounds occupy the middle pocket of the palmitate binding site, while in TEAD3, the compound adopts a binding mode that closely mimicked palmitate. Mimicking the binding mode of palmitate will likely result in stabilization of TEAD and likely improved binding to YAP. But binding to the middle pocket in TEAD2 could be the basis for the higher inhibition of TEAD2 and TEAD4 binding to YAP. This binding mode puts the compound closer to the TEAD•YAP protein-protein interface and may cause shifting of the conformation of amino acids at the interface that are responsible for inhibition of the protein-protein interaction. Allostery is a complex process and further studies using tools capable of exploring the dynamical changes of the protein such as NMR will be required to uncover the exact mechanism of inhibition. Future structure-based optimization for the purpose of disrupting TEAD binding to YAP may need to be focused at the central and upper sub-pockets to enhance potency of compounds.

It is interesting to note that the *K*_I_ values for the compounds were generally high. *K*_I_ is the concentration of compound required to achieve half the reaction rate *k*_inact_. *K*_I_ is not the binding affinity, but a high affinity compound will likely require much lower concentrations to achieve the same reaction rate. For allosteric inhibitors, high affinity binding may not be desirable as this may stabilize the protein and possibly enhance rather than inhibit the protein-protein interaction. This is why covalent inhibitors may hold an advantage over non-covalent compounds for allosteric inhibition, as covalent inhibitors are driven by a binding and a reaction step. It is possible to achieve high second order rate *k*_inact_ / *K*_I_ even with a lower affinity compound with covalent inhibitors. In fact, penicillin and the recently FDA-approved KRAS G12C covalent inhibitors are examples of compounds with low affinity to their target, yet they have superior potency. Focusing on enhancing the reaction rate while enhancing molecular recognition to the pocket to increase specificity of the compound may be the best strategy to develop more potent inhibitors, although more studies will be required to establish this for TEADs.

## METHODS

### Protein Expression and Purification

TEAD1 (residues 209-426) was cloned into pET-28a-TEV vector and transformed into *E. coli* BL21 (DE3) cells for expression. The cells were grown at 37°C to OD_600_ of 0.6 and induced with 0.5 mM isopropyl β-D-1-thiogalactopyranoside (IPTG) overnight at 16°C. The cells were harvested by centrifugation and lysed by microfluidizer in lysis buffer [2 x phosphate buffered saline (PBS), 2 mM β-mercaptoethanol (BME)]. The lysate was clarified by centrifugation and the supernatant was loaded onto a 5-mL HisTrap HP column (Cytiva Life Sciences, Marlborough, MA). The column was washed with 10 column volume (CV) lysis buffer and eluted by linear gradient against elution buffer (2 x PBS, 500 mM imidazole, pH 7.8, 2 mM BME). The eluted protein was concentrated and treated with 6 mM hydroxylamine for 2.5 h at room temperature (rt). The sample was further purified on Superdex 75 pg XK16/60 size-exclusion chromatography (SEC) column (Cytiva Life Sciences, Marlborough, MA) in PBS with 1 mM dithiothreitol (DTT).

TEAD2 (residues 217-447) was cloned into pET-28a-TEV vector and transformed into *E. coli* BL21 (DE3) cells for expression. The cells were grown at 37°C to OD_600_ of 0.6-0.8 and induced with 0.5 mM IPTG overnight at 16°C. The cells were harvested by centrifugation and lysed by microfluidizer in lysis buffer (500 mM NaCl, 50 mM HEPES pH 7.5, 1 mM tris(2-carboxyethyl) phosphine (TCEP)). The lysate was clarified by centrifugation and the supernatant was loaded onto a 1-mL PureCube Ni-INDIGO column (Cube BioTech, Monheim, Germany). The column was washed with 30 CV wash buffer (300 mM NaCl, 25 mM HEPES pH 7.5, 30 mM imidazole, 5% glycerol, 1 mM TCEP) and eluted with 25 mL elution buffer (300 mM imidazole, 300 mM NaCl, 25 mM HEPES pH 7.5, 5% glycerol, 1 mM TCEP). The eluent was concentrated and treated with 6 mM hydroxylamine for 2.5 h at rt. The sample was then purified on Superdex 75pg XK16/60 SEC column (Cytiva Life Sciences, Marlborough, MA) in 100 mM NaCl, 10 mM HEPES pH 7.2, 1 mM TCEP.

For crystallization, the protein eluted from the PureCube Ni-INDIGO column was cleaved with 25:1 w/w tobacco-etch virus (TEV) enzyme for 16-24 h at 4°C while dialyzing against 1 L wash buffer. After cleavage, the reaction was passed through the PureCube Ni-INDIGO column to remove TEV enzyme and uncleaved His-TEV-TEAD2. The sample was treated with hydroxylamine and purified by SEC as above.

TEAD3 (residues 216-435) was cloned into pET-28a-TEV vector and transformed into *E. coli* BL21 (DE3) cells for expression. The cells were grown at 37°C to OD_600_ of 0.6 and induced with 0.5 mM IPTG overnight at 16°C. The cells were harvested by centrifugation and lysed by microfluidizer in lysis buffer (500 mM NaCl, 20 mM NaPi pH 7.8, 2 mM BME). The lysate was clarified by centrifugation and the supernatant was loaded onto a 5-mL HisTrap HP column (Cytiva Life Sciences, Marlborough, MA). The column was washed with 10 CV lysis buffer and eluted by linear gradient against elution buffer (1 M NaCl, 0.5 M imidazole, 2 mM BME, pH 6.5). The eluent was concentrated and treated with 6 mM hydroxylamine for 2.5 h at rt. The sample was further purified on Superdex 75pg XK16/60 SEC column (Cytiva Life Sciences, Marlborough, MA) in PBS with 2 mM DTT.

For crystallization, the protein eluted from the HisTrap HP column was cleaved with 50:1 w/w TEV enzyme for 24 h at 4°C while dialyzing against 1 L buffer containing 500 mM NaCl, 20 mM NaPi pH 7.8, 2 mM DTT. After cleavage, the reaction solution was passed through a PureCube Ni-INDIGO column to remove TEV enzyme and uncleaved His-TEV-TEAD3. The sample was treated with 6 mM hydroxylamine for 2.5 h at rt and purified on a Superdex 75pg XK16/60 SEC column (Cytiva Life Sciences, Marlborough, MA) in 100 mM NaCl, 10 mM HEPES pH 7.2, 1 mM TCEP.

TEAD4 (217–434) and TEAD4^C367S^ mutant were cloned into pGEX-6P-1 vector and transformed into *E. coli* BL21 (DE3) cells for expression. The cells were grown at 37°C to OD_600_ of 0.6 and induced with 0.5 mM IPTG overnight at 16°C. The cells were harvested by centrifugation and lysed by microfluidizer in lysis buffer (200 mM NaCl, 20 mM Tris pH 8.0, 2 mM DTT). The lysate was clarified by centrifugation and the supernatant was loaded onto a 5-mL GSTrap HP column (Cytiva Life Sciences, Marlborough, MA). The column was washed with 20 CV lysis buffer and eluted with 5 CV elution buffer (200 mM NaCl, 20 mM Tris pH 8.0, 2 mM DTT, 10 mM glutathione). The eluted protein was concentrated and purified on Superdex 200pg 26/60 column (Cytiva Life Sciences, Marlborough, MA) in lysis buffer. The protein was cleaved with 50:1 w/w GST-HRV-3C enzyme (ThermoFisher, Waltham, MA) for 24 h at 4°C while dialyzing against 2 x 1 L lysis buffer. After cleavage, the reaction was passed through the GSTrap HP column to remove GST-HRV-3C as well as cleaved GST.

### Fluorescence Polarization Binding Assay

Inhibition of TEAD interaction with YAP was investigated using a fluorescently labeled YAP TEAD-binding domain (residues 60-99) [FAM-DSETDLEALFNAVMNPKTANVPQTVPMCLRKLPASFCKPP]. The cysteines in the peptide are crosslinked. In a 384-well black polystyrene plate (Cat. No. 262260; Nunc, Roskilde, Denmark), 40 µL TEAD in assay buffer (PBS, 0.01% Triton X-100) was added. Compounds were added in a 5 µL mixture of assay buffer and 20% v/v dimethylsulfoxide (DMSO) and incubated for 24 h at 4°C. After the incubation, 5 µL fluorescent peptide was added and the polarization was measured on an Envision Multilabel Plate Reader (PerkinElmer, Waltham, MA) using a filter set with excitation and emission wavelengths of 485 and 535 nm, respectively. The final concentrations for TEAD1-4 were 250, 64, 64 and 64 nM, respectively, while the concentration of the fluorescent peptide was 16 nM for all experiments. Percent inhibition was calculated relative to the maximum polarization control (no compound) and minimum polarization control (no TEAD). For fragment screening, TEAD4 was incubated with 50 µM fragments for 24 h at 4°C. For concentration-dependent analysis, the compounds were first serially diluted in DMSO to keep DMSO concentrations identical between samples.

For time-dependent inhibition of TEAD4 by **14** (ACR-374), multiple identical sample plates were set up at the same time and read after 0.5 h, 6 h, 24 h and 48 h. Inhibition versus time plot was fit with an exponential function 𝑃𝑒𝑟𝑐𝑒𝑛𝑡 𝐼𝑛ℎ𝑖𝑏𝑖𝑡𝑖𝑜𝑛 = 𝐸𝑥𝑡𝑒𝑛𝑡 × (1 − 𝑒^-k_obs_×Time^) to determine pseudo first-order rate constant *k*_obs_. The *k*_obs_ values were plotted against compound concentrations and fit with a hyperbolic function 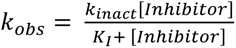 to determine maximum rate of adduct formation *k*_inact_ and the binding constant *K*_I_.

### Protein Mass Spectrometry

TEAD1-4 and TEAD4^C367S^ at 2.5 µM concentration were incubated with 100 µM acrylamide compounds for 24 h at 4°C in PBS. After the incubation, the samples were centrifuged at 21,000 x *g* to remove precipitants and dust. The samples were injected into Agilent 1200 liquid chromatography system (Agilent, Santa Clara, CA) equipped with Zorbax 300-SB C3 column (Agilent, Santa Clara, CA). Protein sample was separated from buffer salts on the column and detected on Agilent 6520 Accurate Mass Q-TOF (Agilent, Santa Clara, CA).

For the determination of the *k*_inact_/*K*_I_, 2.5 µM TEAD1-4 were incubated with varying concentrations (3.1-100 µM) of compounds at 4 °C. After varying amounts of time (30 s to 24 h) aliquots were taken from each sample and quenched with 0.1 M formic acid. The samples were then tested as above by mass spectrometry. The increase in TEAD-compound adduct peak relative to the total TEAD peaks was plotted against time for all concentrations of the compounds, and the observed rate constant of adduct formation (*k*_obs_) was determined by fitting an exponential function. The *k*_obs_ values were then plotted against their respective compound concentrations, and a hyperbolic function 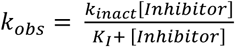 was fitted to determine the *k*_inact_ and *K* values.

### Crystallography

TEAD2 (4-5 mg/mL) crystals were grown in HEPES pH 7.2-7.4, sodium formate 1.8-2.6 M at 20 °C using the hanging-drop vapor-diffusion method. TEAD2-compound complexes were obtained by soaking the crystals for 1-3 days in reservoir solution supplemented with 2-5 mM compound. Crystals were harvested, cryo-protected in reservoir solution supplemented with 25% glycerol and flash-cooled in liquid nitrogen.

TEAD3 (10-12 mg/mL) was mixed with 2-5 mM **14** (ACR-374) and incubated for 2 hours. Drops were prepared by mixing the complex with reservoir solution in 1:1 (v/v) relationship. Crystals grew after several weeks in 0.1 M sodium acetate pH 5.0 – 5.5, 0.1 M calcium acetate and 4-8% PEG4000 at 20°C using the sitting-drop vapor-diffusion method. Crystals were harvested, cryo-protected in reservoir solution supplemented with 30% glycerol and flash-cooled in liquid nitrogen.

Diffraction data were collected at 100 K at the Beamline station 4.2.2 at the Advanced Light Source (Berkeley National Laboratory, CA) and were initially indexed, integrated, and scaled using XDS (42). All data sets presented severe anisotropy, with useful data extending to modest resolution (see **Table S2**). Thus, data sets were re-processed, and anisotropy analysis was performed using the STARANISO server (43). Anisotropic completeness was obtained by fitting an ellipsoid using least squares to the resulting cut-off surface points. Cut-off criteria were Rpim < 0.6, I/σ > 1.2 and CC1/2 > 0.5. Final data sets, after anisotropic correction, were used to perform molecular replacement using PHASER and PDB code 6E5G (17) for TEAD2 and 5EMW for TEAD3 as search models. Successive cycles of automatic building in Autobuild (PHENIX) and manual building in Coot, as well as refinement (PHENIX Refine (44)) led to complete models. MolProbity software (45) was used to assess the geometric quality of the models, and PyMoL (46) was used to generate molecular images. Data collection and refinement statistics are indicated (**Table S2**).

## ACKNOWLEDGMENTS

The research was supported by an American Cancer Society Research Scholar Grant RSG-12-092-01-CDD (SOM), a Vera Bradley Foundation fellowship (KB), a Vera Bradley Foundation grant (SOM), an Indiana University Simon Cancer Center Near Miss Initiative grant (SOM), and the 100 Voices of Hope (SOM). The research was supported by the Department of Veteran Affairs (I01BX005188) [SOM], and the National Institutes of Health (R01CA264471) [SOM].

## AUTHOR CONTRIBUTIONS

K. B., M. K. G., M. J. Z. carried out biochemical studies. G. G-G. grew crystals and collected data, and K. B. and G. G-G. solved crystal structures. K. B., G. G-G., and S. O. M. designed experiments and analyzed data. S. O. M. designed compound structures. K. B., G. G-G., and S. O. M. wrote the paper.

## DECLARATION OF INTERESTS

None

## ABBREVIATIONS

TEAD: transcriptional enhanced associate domain
YAP: Yes-associated protein
FP: fluorescence polarization
PBS: phosphate buffered saline
BME: β-mercaptoethanol
SEC: size exclusion chromatography
DTT: dithiothreitol
TCEP: tris(2-carboxyethyl)phosphine
TEV: tobacco etch virus
GST: glutathione-S-transferase
HRV: human rhinovirus
DMSO: dimethylsulfoxide
Q-TOF: quadrupole time-of-flight

## Notes

### Competing Interest Statement

The authors have declared no competing interest.

## REFERENCES

1. Zhao, B., Li, L., Tumaneng, K., Wang, C. Y., and Guan, K. L. (2010) A coordinated phosphorylation by Lats and CK1 regulates YAP stability through SCF(beta-TRCP) Genes Dev 24, 72–85

2. Liu, C. Y., Zha, Z. Y., Zhou, X., Zhang, H., Huang, W., Zhao, D., et al. (2010) The hippo tumor pathway promotes TAZ degradation by phosphorylating a phosphodegron and recruiting the SCF{beta}-TrCP E3 ligase J Biol Chem 285, 37159–37169

3. Kim, Y., and Jho, E. H. (2018) Regulation of the Hippo signaling pathway by ubiquitin modification BMB Rep 51, 143–150

4. Zhao, B., Ye, X., Yu, J., Li, L., Li, W., Li, S. et al. (2008) TEAD mediates YAP-dependent gene induction and growth control Genes & Development 22, 1962–1971

5. Zhang, J., Smolen, G. A., and Haber, D. A. (2008) Negative Regulation of YAP by LATS1 Underscores Evolutionary Conservation of the Drosophila Hippo Pathway Cancer Research 68, 2789–2794

6. Hyun, Young, Yu, B., Moroishi, T., Mo, J.-S., Steven et al. (2015) Alternative Wnt Signaling Activates YAP/TAZ Cell 162, 780–794

7. Seo, E., Basu-Roy, U., Gunaratne, Preethi H., Coarfa, C., Lim, D.-S., Basilico, C. et al. (2013) SOX2 Regulates YAP1 to Maintain Stemness and Determine Cell Fate in the Osteo-Adipo Lineage Cell Reports 3, 2075–2087

8. Lee, D.-H., Park, J. O., Kim, T.-S., Kim, S.-K., Kim, T.-H., Kim, M.-C., et al. (2016) LATS-YAP/TAZ controls lineage specification by regulating TGFβ signaling and Hnf4α expression during liver development Nature Communications 7, 11961

9. Lai, D., and Yang, X. (2013) BMP4 is a novel transcriptional target and mediator of mammary cell migration downstream of the Hippo pathway component TAZ Cellular Signalling 25, 1720–1728

10. Zhang, J., Ji, J.-Y., Yu, M., Overholtzer, M., Smolen, G. A., Wang, R. et al. (2009) YAP-dependent induction of amphiregulin identifies a non-cell-autonomous component of the Hippo pathway Nature Cell Biology 11, 1444–1450

11. Song, S., Honjo, S., Jin, J., Chang, S.-S., Scott, A. W., Chen, Q. et al. (2015) The Hippo Coactivator YAP1 Mediates EGFR Overexpression and Confers Chemoresistance in Esophageal Cancer Clinical Cancer Research 21, 2580–2590

12. Feng, J., Yang, H., Zhang, Y., Wei, H., Zhu, Z., Zhu, B. et al. (2017) Tumor cell-derived lactate induces TAZ-dependent upregulation of PD-L1 through GPR81 in human lung cancer cells Oncogene 36, 5829–5839

13. Kim, M. H., Kim, C. G., Kim, S. K., Shin, S. J., Choe, E. A., Park, S. H. et al. (2018) YAP-Induced PD-L1 Expression Drives Immune Evasion in BRAFi-Resistant Melanoma Cancer Immunol Res 6, 255–266

14. Miao, J., Hsu, P. C., Yang, Y. L., Xu, Z., Dai, Y., Wang, Y., et al. (2017) YAP regulates PD-L1 expression in human NSCLC cells Oncotarget 8, 114576–114587

15. Neto-Silva, R. M., de Beco, S., and Johnston, L. A. (2010) Evidence for a growth-stabilizing regulatory feedback mechanism between Myc and Yorkie, the Drosophila homolog of Yap Dev Cell 19, 507–520

16. Rajbhandari, P., Lopez, G., Capdevila, C., Salvatori, B., Yu, J., Rodriguez-Barrueco, R. et al. (2018) Cross-Cohort Analysis Identifies a TEAD4-MYCN Positive Feedback Loop as the Core Regulatory Element of High-Risk Neuroblastoma Cancer Discov 8, 582–599

17. Bum-Erdene, K., Zhou, D., Gonzalez-Gutierrez, G., Ghozayel, M. K., Si, Y., Xu, D. et al. (2019) Small-Molecule Covalent Modification of Conserved Cysteine Leads to Allosteric Inhibition of the TEADYap Protein-Protein Interaction Cell Chem Biol 26, 378–389 e313

18. Mesrouze, Y., Meyerhofer, M., Bokhovchuk, F., Fontana, P., Zimmermann, C., Martin, T. et al. (2017) Effect of the acylation of TEAD4 on its interaction with co-activators YAP and TAZ Protein Sci 26, 2399–2409

19. Li, Z., Zhao, B., Wang, P., Chen, F., Dong, Z., Yang, H. et al. (2010) Structural insights into the YAP and TEAD complex Genes & development 24, 235–240

20. Tian, W., Yu, J., Tomchick, D. R., Pan, D., and Luo, X. (2010) Structural and functional analysis of the YAP-binding domain of human TEAD2 Proc Natl Acad Sci U S A 107, 7293–7298

21. Noland, Cameron L., Gierke, S., Schnier, Paul D., Murray, J., Sandoval, Wendy N., Sagolla, M. et al. (2016) Palmitoylation of TEAD Transcription Factors Is Required for Their Stability and Function in Hippo Pathway Signaling Structure 24, 179–186

22. Chan, P., Han, X., Zheng, B., DeRan, M., Yu, J., Jarugumilli, G. K., et al. (2016) Autopalmitoylation of TEAD proteins regulates transcriptional output of the Hippo pathway *Nature chemical biology*

23. Holden, J. K., Crawford, J. J., Noland, C. L., Schmidt, S., Zbieg, J. R., Lacap, J. A., et al. (2020) Small Molecule Dysregulation of TEAD Lipidation Induces a Dominant-Negative Inhibition of Hippo Pathway Signaling Cell Reports 31, 107809

24. Bum-Erdene, K., Yeh, I. J., Gonzalez-Gutierrez, G., Ghozayel, M. K., Pollok, K., and Meroueh, S. O. (2023) Small-Molecule Cyanamide Pan-TEAD.YAP1 Covalent Antagonists J Med Chem 66, 266–284

25. Lu, W., Wang, J., Li, Y., Tao, H., Xiong, H., Lian, F. et al. (2019) Discovery and biological evaluation of vinylsulfonamide derivatives as highly potent, covalent TEAD autopalmitoylation inhibitors European Journal of Medicinal Chemistry 184, 111767

26. Lu, T., Li, Y., Lu, W., Spitters, T., Fang, X., Wang, J. et al. (2021) Discovery of a subtype-selective, covalent inhibitor against palmitoylation pocket of TEAD3 Acta Pharmaceutica Sinica B 11, 3206–3219

27. Kaneda, A., Seike, T., Danjo, T., Nakajima, T., Otsubo, N., Yamaguchi, D. et al. (2020) The novel potent TEAD inhibitor, K-975, inhibits YAP1/TAZ-TEAD protein-protein interactions and exerts an anti-tumor effect on malignant pleural mesothelioma Am J Cancer Res 10, 4399–4415

28. Karatas, H., Akbarzadeh, M., Adihou, H., Hahne, G., Pobbati, A. V., Yihui Ng, E. et al. (2020) Discovery of Covalent Inhibitors Targeting the Transcriptional Enhanced Associate Domain Central Pocket Journal of Medicinal Chemistry 63, 11972–11989

29. Kurppa, K. J., Liu, Y., To, C., Zhang, T., Fan, M., Vajdi, A. et al. (2020) Treatment-Induced Tumor Dormancy through YAP-Mediated Transcriptional Reprogramming of the Apoptotic Pathway Cancer Cell 37, 104–122.e112

30. Fan, M., Lu, W., Che, J., Kwiatkowski, N. P., Gao, Y., Seo, H.-S., et al. (2022) Covalent disruptor of YAP-TEAD association suppresses defective Hippo signaling eLife 11, e78810

31. Lu, W., Fan, M., Ji, W., Tse, J., You, I., Ficarro, S. B. et al. (2023) Structure-Based Design of Y-Shaped Covalent TEAD Inhibitors Journal of Medicinal Chemistry 66, 4617–4632

32. Nutsch, K., Song, L., Chen, E., Hull, M., Chatterjee, A. K., Chen, J. J. et al. (2023) A covalent inhibitor of the YAP-TEAD transcriptional complex identified by high-throughput screening RSC Chem Biol 4, 894–905

33. Hillen, H., Candi, A., Vanderhoydonck, B., Kowalczyk, W., Sansores-Garcia, L., Kesikiadou, E. C. et al. (2023) A novel irreversible TEAD inhibitor, SWTX-143, blocks Hippo pathway transcriptional output and causes tumor regression in preclinical mesothelioma models Mol Cancer Ther 10.1158/1535-7163.MCT-22-0681

34. Li, Q., Sun, Y., Jarugumilli, G. K., Liu, S., Dang, K., Cotton, J. L. et al. (2020) Lats1/2 Sustain Intestinal Stem Cells and Wnt Activation through TEAD-Dependent and Independent Transcription Cell Stem Cell 26, 675–692.e678

35. Sun, Y., Hu, L., Tao, Z., Jarugumilli, G. K., Erb, H., Singh, A. et al. (2022) Pharmacological blockade of TEAD–YAP reveals its therapeutic limitation in cancer cells Nature Communications 13, 6744

36. Li, L., Li, R., and Wang, Y. (2022) Identification of Small-molecule YAP-TEAD inhibitors by High-throughput docking for the Treatment of colorectal cancer Bioorganic Chemistry 122, 105707

37. Tang, T. T., Konradi, A. W., Feng, Y., Peng, X., Ma, M., Li, J. et al. (2021) Small Molecule Inhibitors of TEAD Auto-palmitoylation Selectively Inhibit Proliferation and Tumor Growth of NF2-deficient Mesothelioma Molecular Cancer Therapeutics 20, 986–998

38. Hu, L., Sun, Y., Liu, S., Erb, H., Singh, A., Mao, J. et al. (2022) Discovery of a new class of reversible TEA domain transcription factor inhibitors with a novel binding mode *eLife* **11**, e80210

39. Hagenbeek, T. J., Zbieg, J. R., Hafner, M., Mroue, R., Lacap, J. A., Sodir, N. M. et al. (2023) An allosteric pan-TEAD inhibitor blocks oncogenic YAP/TAZ signaling and overcomes KRAS G12C inhibitor resistance Nature Cancer 4, 812–828

40. Heinrich, T., Peterson, C., Schneider, R., Garg, S., Schwarz, D., Gunera, J. et al. (2022) Optimization of TEAD P-Site Binding Fragment Hit into In Vivo Active Lead MSC-4106 Journal of Medicinal Chemistry 65, 9206–9229

41. Geoghegan, K. F., Dixon, H. B., Rosner, P. J., Hoth, L. R., Lanzetti, A. J., Borzilleri, K. A. et al. (1999) Spontaneous alpha-N-6-phosphogluconoylation of a “His tag” in Escherichia coli: the cause of extra mass of 258 or 178 Da in fusion proteins Anal Biochem 267, 169–184

42. Kabsch, W. (2010) Xds Acta Crystallogr D Biol Crystallogr 66, 125–132

43. Tickle, I. J., Flensburg, C., Keller, P., Paciorek, W., Sharff, A., Vonrhein, C. et al. (2018) STARANISO Global Phasing Ltd., Cambridge, UK

44. Liebschner, D., Afonine, P. V., Baker, M. L., Bunkoczi, G., Chen, V. B., Croll, T. I. et al. (2019) Macromolecular structure determination using X-rays, neutrons and electrons: recent developments in Phenix Acta Crystallogr D Struct Biol 75, 861–877

45. Williams, C. J., Headd, J. J., Moriarty, N. W., Prisant, M. G., Videau, L. L., Deis, L. N. et al. (2018) MolProbity: More and better reference data for improved all-atom structure validation Protein Sci 27, 293–315

46. LLC, S., and Delano, W. (2020) PyMOL

