## Supporting Information for "A Fragment Screen Identifies Acrylamide Covalent Inhibitors of the TEAD•YAP Protein-Protein Interaction"

**CONTENTS**

**Supporting Figure S1.** Chemical Structures of Fragment Screen Hits

**Supporting Figure S2.** Characterization of Fragment Screen Hits

**Supporting Figure S3.** Reaction of **1** (ACR-021) Derivatives to TEADs

**Supporting Figure S4**. Time-Dependent Mass Spectrometry

**Supporting Figure S5**. Crystal Structure of TEAD3 in Complex with **14** (ACR-374)

**Supporting Figure S6.** The Chemical Structures of TEAD Lipid Pocket Inhibitors

**Supporting Figure S7.** Inhibition of TEAD1-4 binding to YAP1 by K-975

**Supporting Figure S8.** HPLC Data for Synthesized Acrylamides

**Supporting Table S1.** Fragment Hit Inhibition and Reaction

**Supporting Table S2.** X-Ray Crystal Diffraction Data Collection and Refinement Statistics

**Chemical Synthesis**

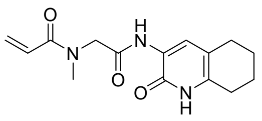

**3** ( ACR-047)

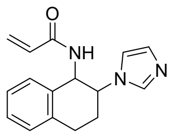

**4** ( ACR-057)

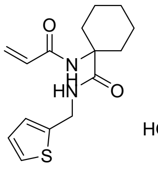

**5** ( ACR-083)

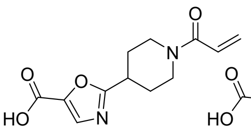

**6** ( ACR-130)

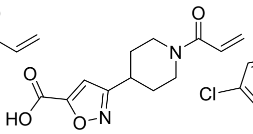

**7** ( ACR-147)

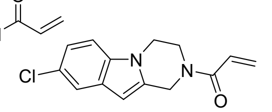

**8** ( ACR-184)

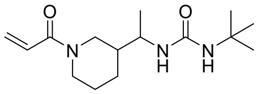

**9** ( ACR-195)

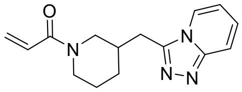

**10** ( ACR-199)

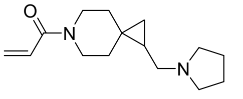

**11** ( ACR-300)

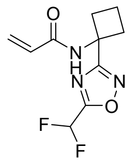

**12** ( ACR-338)

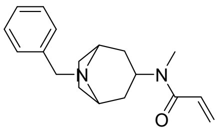

**13** ( ACR-345)

**Supporting Figure S1.** Chemical Structures of Fragment Screen Hits

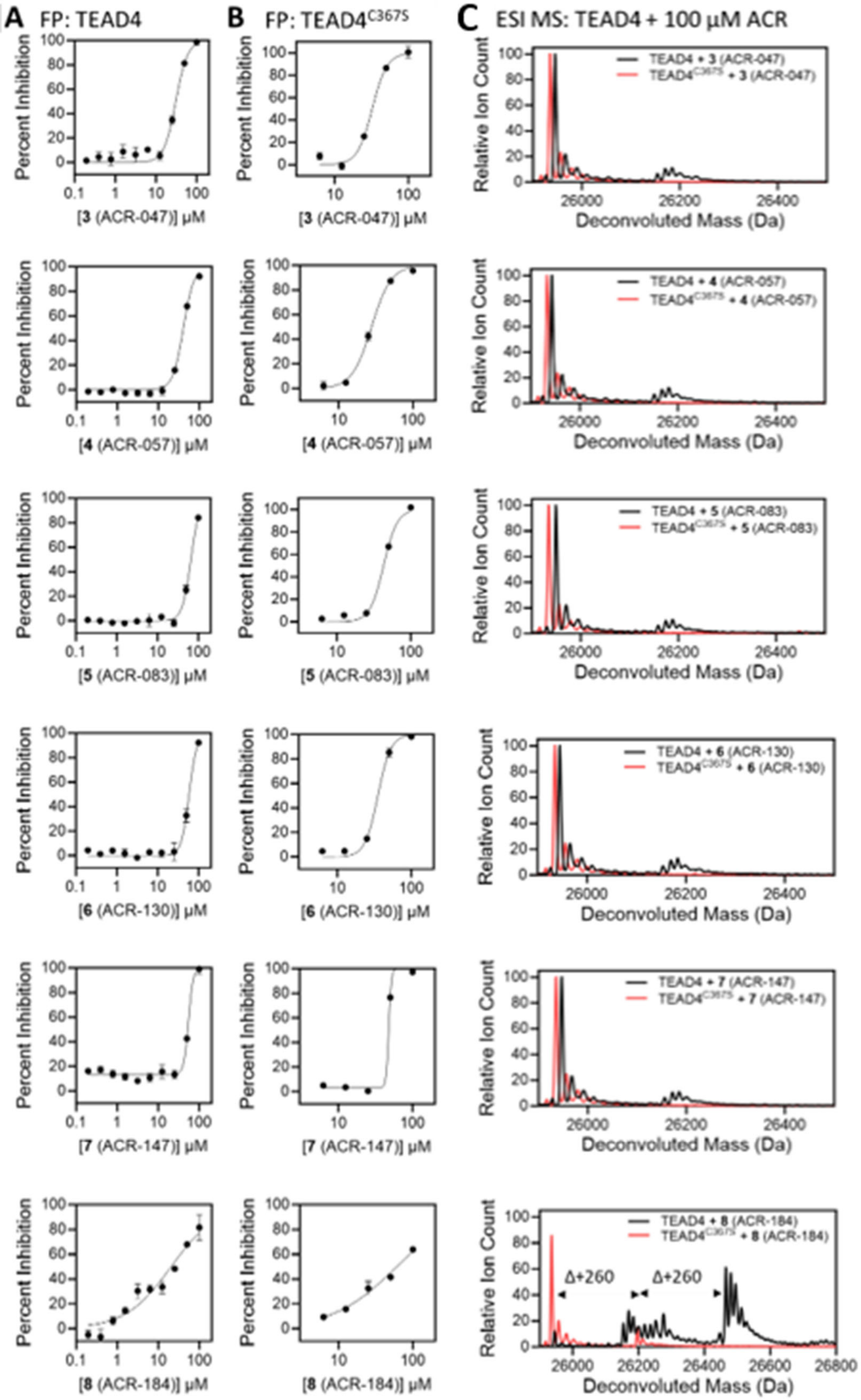

A FP: TEAD4
B FP: TEAD4C367S
C ESI MS: TEAD4 + 100 µM ACR
Percent Inhibition
Relative Ion Count
Deconvoluted Mass (Da)
[3 (ACR-047)] µM
[4 (ACR-057)] µM
[5 (ACR-083)] µM
[6 (ACR-130)] µM
[7 (ACR-147)] µM
[8 (ACR-184)] µM
TEAD4 + 3 (ACR-047)
TEAD4C367S + 3 (ACR-047)
TEAD4 + 4 (ACR-057)
TEAD4C367S + 4 (ACR-057)
TEAD4 + 5 (ACR-083)
TEAD4C367S + 5 (ACR-083)
TEAD4 + 6 (ACR-130)
TEAD4C367S + 6 (ACR-130)
TEAD4 + 7 (ACR-147)
TEAD4C367S + 7 (ACR-147)
TEAD4 + 8 (ACR-184)
TEAD4C367S + 8 (ACR-184)
Δ+260
Δ+260

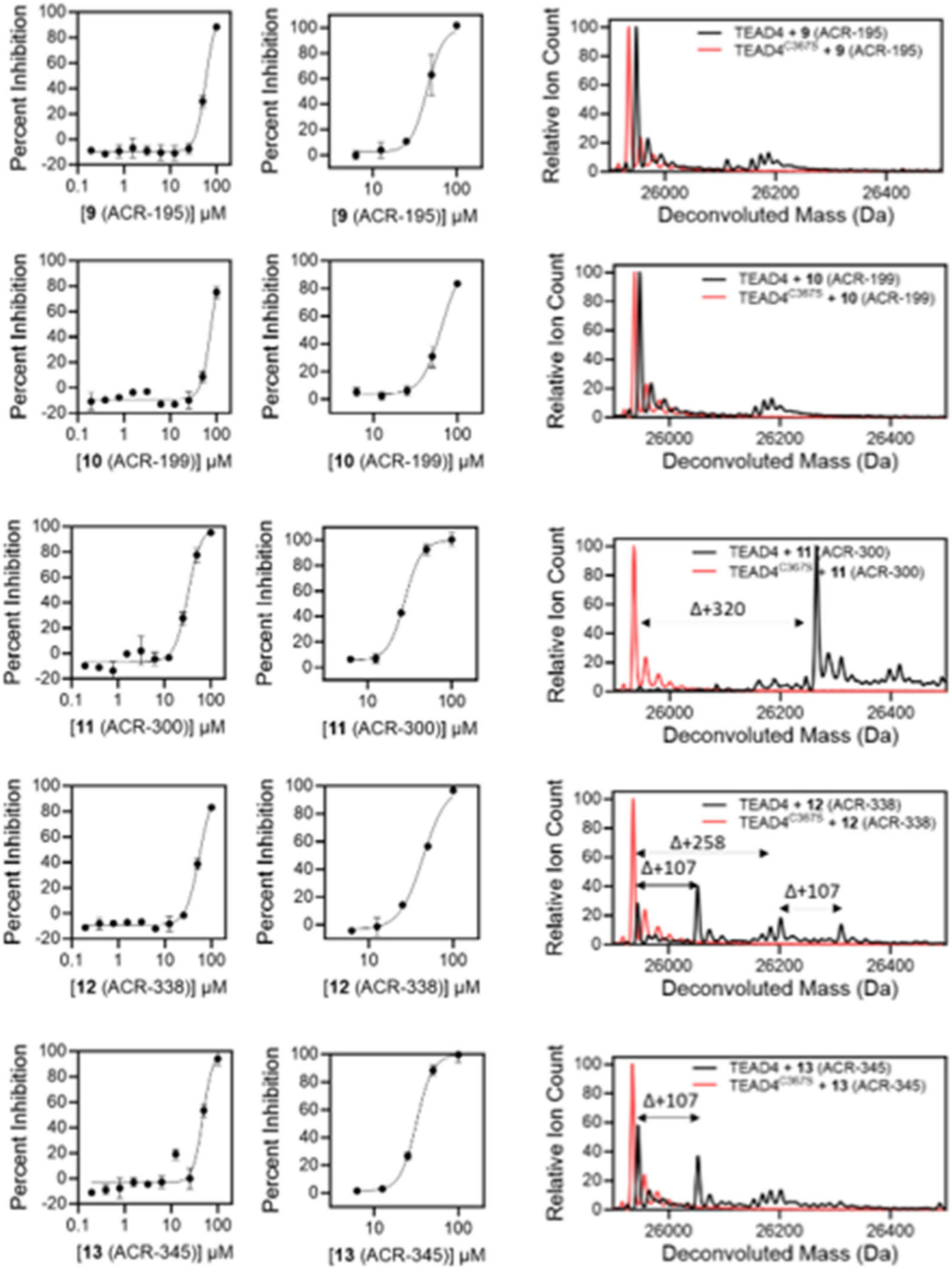


**Supporting Figure S2.** Characterization of Fragment Screen Hits.

A set of 13 hit compounds were found to inhibit TEAD4•YAP1 binding. The compounds were tested for inhibition of **(A)** TEAD4 wildtype and **(B)** TEAD4$^{C367S}$ mutant in a concentration-dependent manner after 24 h incubation at 4°C. **(C)** Compound reaction to TEAD4 and TEAD4$^{C367S}$ was detected by whole-protein mass spectrometry after 24 h incubation at 4°C.

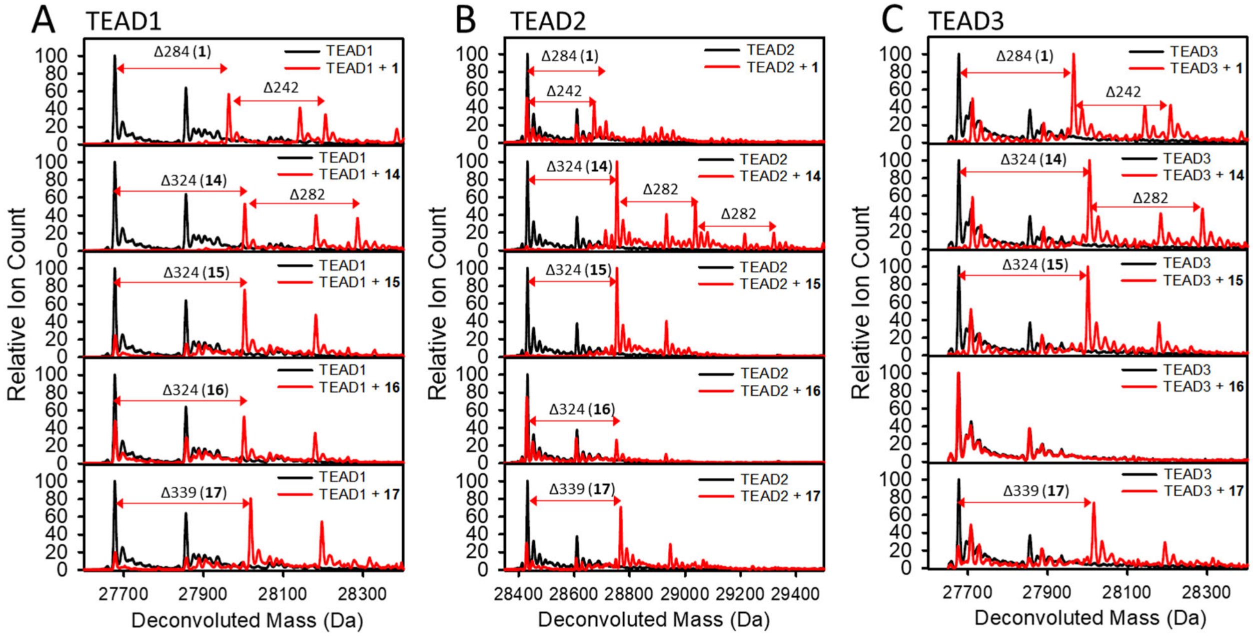


**Supporting Figure S3.** Reaction of **1** (ACR-021) Derivatives to TEADs. Whole-protein mass spectrometry of TEAD1, TEAD2 and TEAD3 after 24 h 4°C incubation with 100 µM compound. The extent of adduct formation with TEAD1-3 for each compound is listed in **Table 1**. **(A)** TEAD1 was detected with two peaks at 27679 Da and 27857 Da. The second peak at 27857 Da corresponds to N-terminal gluconoylation of TEAD1. Fragment adducts to TEAD1 were detected at mass differences equal to the molecular weight of the compound. Fragment **1** (ACR-021) and **15** (ACR-374) formed additional adducts with masses that are 42 less than the molecular weight of the compound. **(B)** TEAD2 was detected with two peaks at 28432 Da and 28610 Da. The second peak at 28610 Da corresponds to N-terminal gluconoylation of TEAD2. Fragment adducts to TEAD2 were detected at mass differences equal to the molecular weight of the compound. Fragment **1** (ACR-021) and **15** (ACR-374) formed additional adducts with masses that are 42 less than the molecular weight of the compound. **(C)** TEAD3 was detected with two peaks at 27682 Da and 27860 Da. The second peak at 27860 Da corresponds to N-terminal gluconoylation of TEAD3. Fragment adducts to TEAD3 were detected at mass differences equal to the molecular weight of the compound. Fragment **17** (ACR-380) did not form an adduct to TEAD3. Fragment **1** (ACR-021) and **15** (ACR-374) formed additional adducts with masses that are 42 less than the molecular weight of the compound.

TEAD1

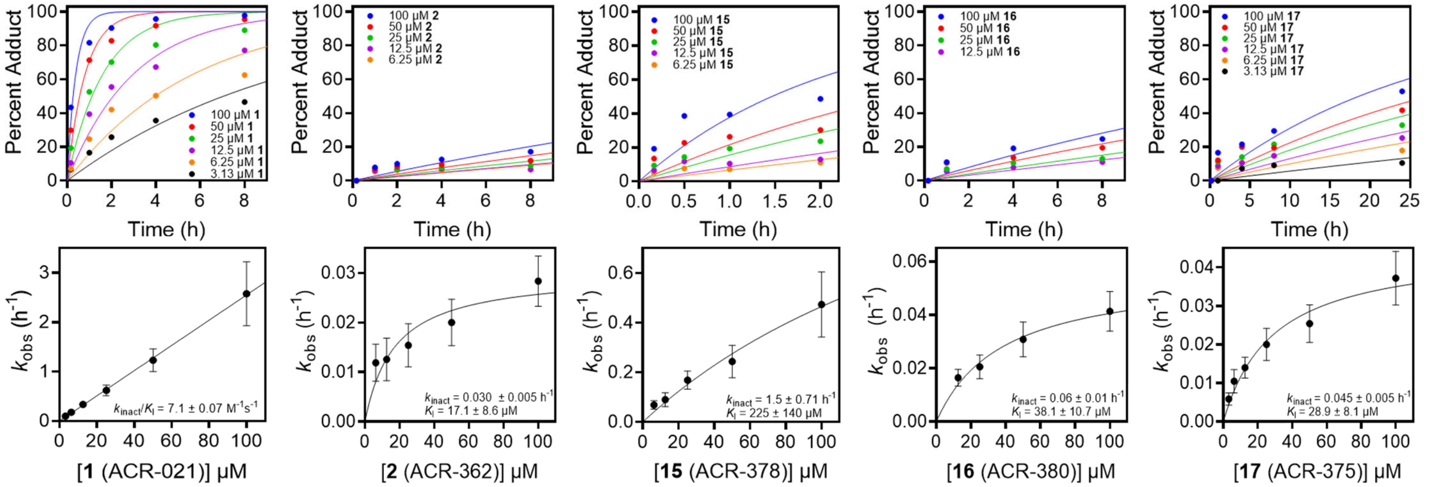

Percent Adduct
Time (h)
100 μM 1
50 μM 1
25 μM 1
12.5 μM 1
6.25 μM 1
3.13 μM 1
100 μM 2
50 μM 2
25 μM 2
12.5 μM 2
6.25 μM 2
100 μM 15
50 μM 15
25 μM 15
12.5 μM 15
6.25 μM 15
100 μM 16
50 μM 16
25 μM 16
12.5 μM 16
100 μM 17
50 μM 17
25 μM 17
12.5 μM 17
6.25 μM 17
3.13 μM 17
$k_{obs}$ ($h^{-1}$)
$k_{inact}/K_I$ = 7.1 ± 0.07 $M^{-1}s^{-1}$
$k_{inact}$ = 0.030 ± 0.005 $h^{-1}$
$K_I$ = 17.1 ± 8.6 μM
$k_{inact}$ = 1.5 ± 0.71 $h^{-1}$
$K_I$ = 225 ± 140 μM
$k_{inact}$ = 0.06 ± 0.01 $h^{-1}$
$K_I$ = 38.1 ± 10.7 μM
$k_{inact}$ = 0.045 ± 0.005 $h^{-1}$
$K_I$ = 28.9 ± 8.1 μM
[1 (ACR-021)] μM
[2 (ACR-362)] μM
[15 (ACR-378)] μM
[16 (ACR-380)] μM
[17 (ACR-375)] μM


TEAD2

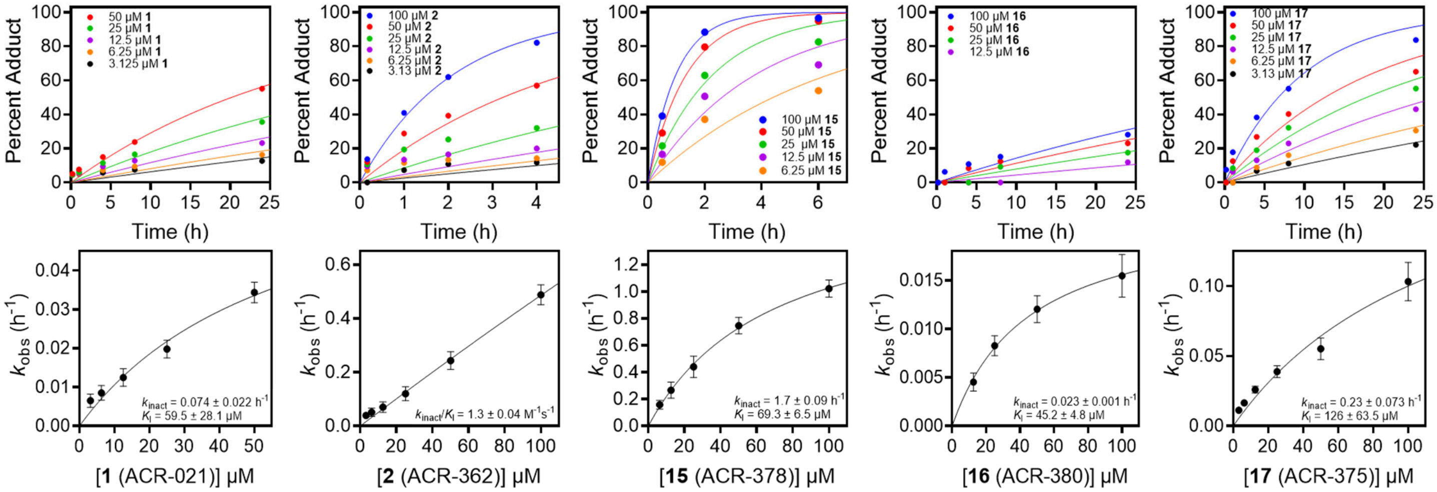

Percent Adduct
Time (h)
50 μM 1
25 μM 1
12.5 μM 1
6.25 μM 1
3.125 μM 1
100 μM 2
50 μM 2
25 μM 2
12.5 μM 2
6.25 μM 2
3.13 μM 2
100 μM 15
50 μM 15
25 μM 15
12.5 μM 15
6.25 μM 15
100 μM 16
50 μM 16
25 μM 16
12.5 μM 16
100 μM 17
50 μM 17
25 μM 17
12.5 μM 17
6.25 μM 17
3.13 μM 17
$k_{obs}$ ($h^{-1}$)
$k_{inact}$ = 0.074 ± 0.022 $h^{-1}$
$K_I$ = 59.5 ± 28.1 μM
$k_{inact}/K_I$ = 1.3 ± 0.04 $M^{-1}s^{-1}$
$k_{inact}$ = 1.7 ± 0.09 $h^{-1}$
$K_I$ = 69.3 ± 6.5 μM
$k_{inact}$ = 0.023 ± 0.001 $h^{-1}$
$K_I$ = 45.2 ± 4.8 μM
$k_{inact}$ = 0.23 ± 0.073 $h^{-1}$
$K_I$ = 126 ± 63.5 μM
[1 (ACR-021)] μM
[2 (ACR-362)] μM
[15 (ACR-378)] μM
[16 (ACR-380)] μM
[17 (ACR-375)] μM

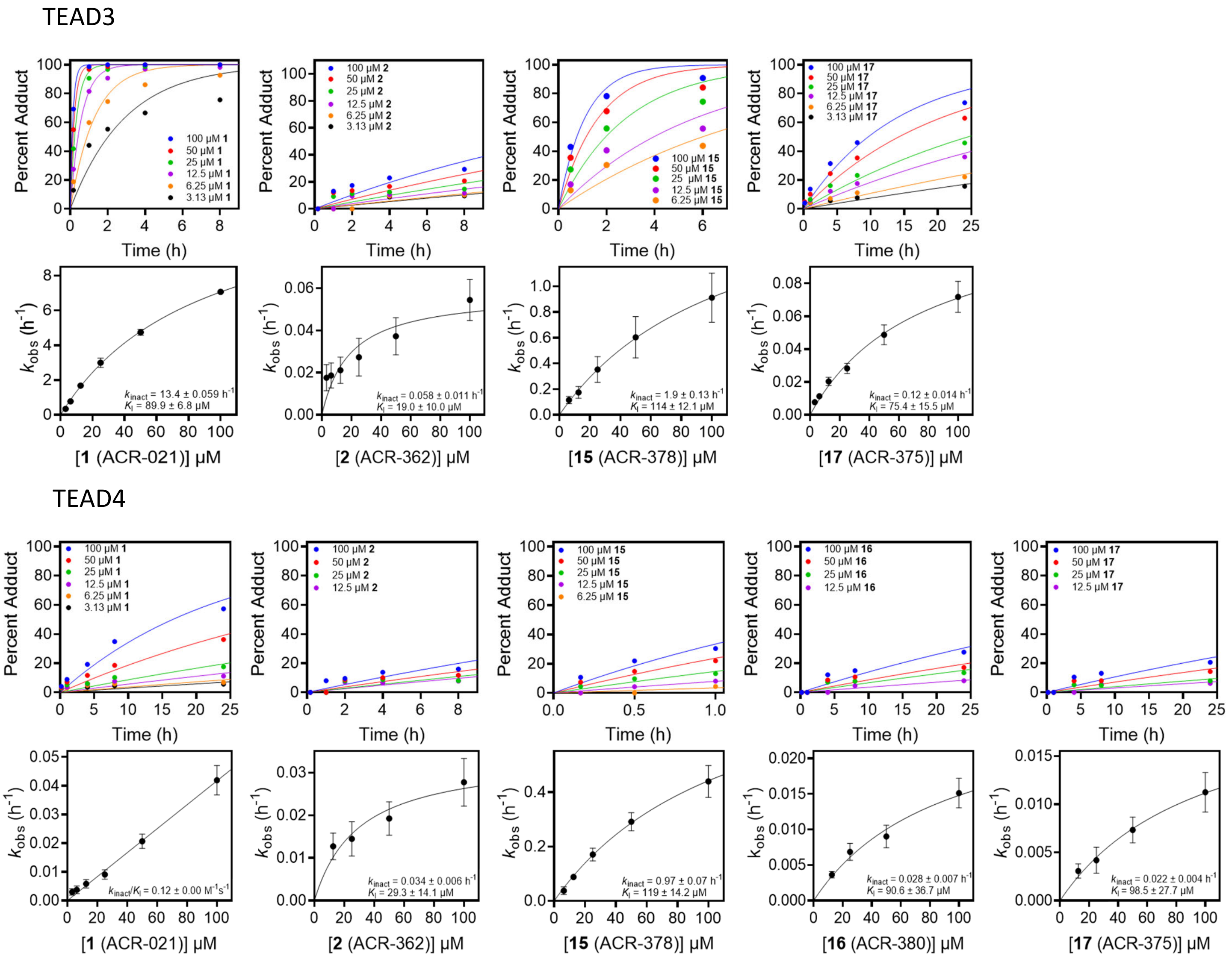


**Supporting Figure S4**. Time-Dependent Mass Spectrometry. TEAD1-4 were incubated with varying concentrations of **1** (ACR-021), **2** (ACR-362), **15** (ACR-378), **16** (ACR-380), and **17** (ACR-375) at 4°C. At indicated time intervals, an aliquot was quenched with 0.1 M formic acid to stop the reaction. The samples were analyzed by whole protein mass spectrometry and percent adduct was determined by the ratio of the adduct intensity to the sum of the protein and adduct peaks. Percent adduct versus time plots were fitted with exponential function $Percent\ Adduct = 100 \times (1 - e^{-k_{obs} \times Time})$ to determine $k_{obs}$. The pseudo first-order rate constant $k_{obs}$ were plotted against their respective concentrations of compounds and fitted with a hyperbolic function $k_{obs} = \frac{k_{inact}[Inhibitor]}{K_I + [Inhibitor]}$ to determine $k_{inact}$ and $K_I$ values.

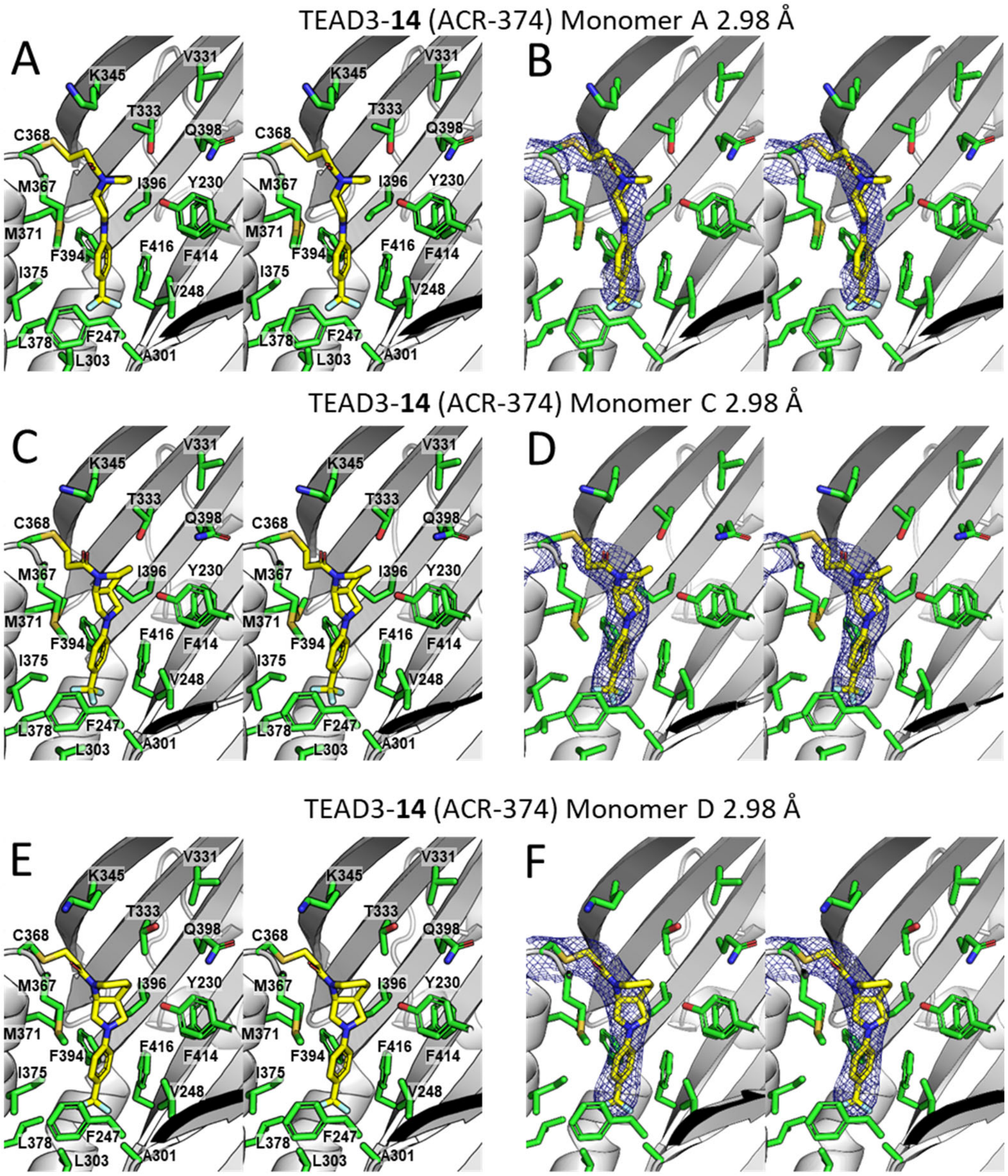


**Supporting Figure S5**. Crystal Structure of TEAD3 in Complex with **14** (ACR-374). **(A)** Stereoview of the covalent binding mode of **14** (ACR-374) [capped-sticks representation with carbon, nitrogen, oxygen, and fluorine in yellow, blue, red, and cyan, respectively] in complex with TEAD3 in monomer A (gray ribbon representation). TEAD3 residues in the pocket are shown in a capped-sticks representation (carbon, nitrogen, oxygen, and sulfur in green, blue, red, and gold, respectively). **(B)** Stereoview of the composite omit map around TEAD3 Cys-368 and **14** (ACR-374) in monomer A. The map is shown in blue mesh at σ 1.0. **(C)** Stereoview of the covalent binding mode of **14** (ACR-374) in complex with TEAD3 in monomer C. Represented as in **A**. **(D)** Stereoview of the composite omit map around TEAD3 Cys-368 and **14** (ACR-374) in monomer C. **(E)** Stereoview of the covalent binding mode of **14** (ACR-374) in complex with TEAD3 in monomer D. Represented as in **A**. **(F)** Stereoview of the composite omit map around TEAD3 Cys-368 and **14** (ACR-374) in monomer D.

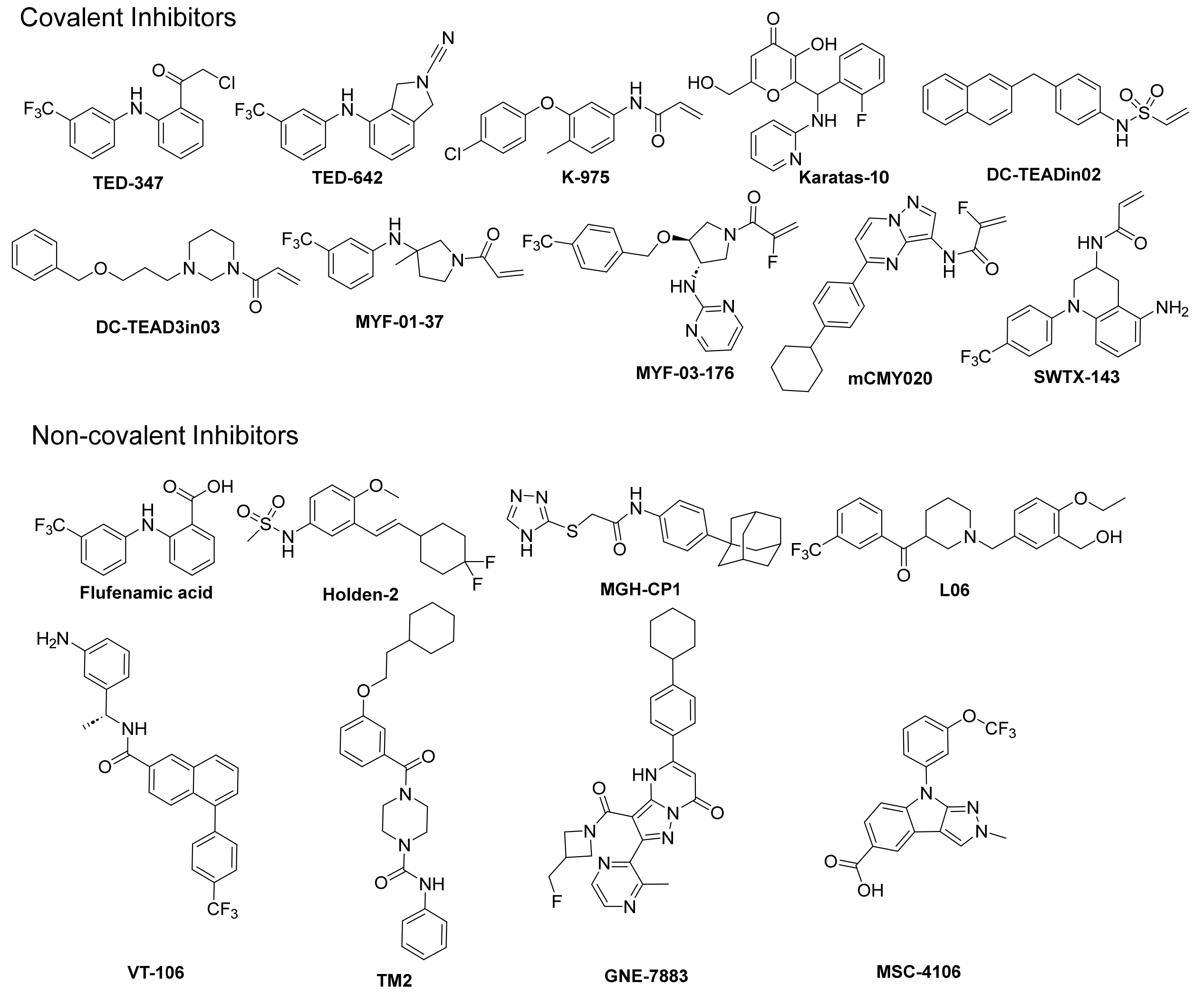


**Supporting Figure S6.** The Chemical Structures of Covalent and Non-Covalent Inhibitors of TEAD Palmitate Pocket. Covalent Inhibitors: TED-347 [1], TED-642 [2], K-975 [3], Karatas-10 [4], DC-TEADin02 [5], DC-TEAD3in03 [6], MYF-01-37 [7], MYF-03-176 [8], mCMY020 [9], SWTX-143 [10]. Non-covalent Inhibitors: Flufenamic acid [11], Holden-2 [12], MGH-CP1 [13], L06 [14], VT-106 [15], TM2 [16], GNE-7883 [17], MSC-4106 [18].

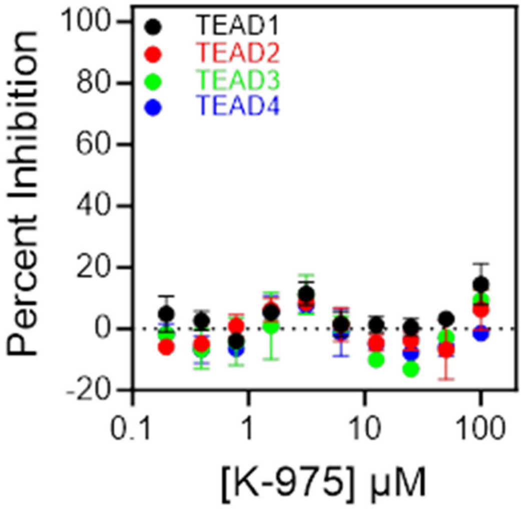


**Supporting Figure S7.** Inhibition of TEAD1-4 binding to YAP1 by K-975. TEAD1-4 were incubated with 0.2-100 μM K-975 for 24 h at 4°C prior to detection of binding to fluorescently labeled YAP1 peptide.

**Supporting Table S1.** Fragment Hit Inhibition and Reaction

| | FP $IC_{50}$ at 24 h 4°C (μM) | | MS Adduct 100 μM 24 h 4°C | |
|---|---|---|---|---|
| **Compound** | **TEAD4 WT** | **TEAD4$^{C367S}$** | **TEAD4** | **TEAD4$^{C367S}$** |
| **3** (ACR-047) | 30.7 ± 2.0 | 32.2 ± 1.7 | 0 | 0 |
| **4** (ACR-057) | 40.7 ± 1.1 | 27.7 ± 0.7 | 0 | 0 |
| **5** (ACR-083) | 65.9 ± 1.5 | 42.9 ± 1.8 | 0 | 0 |
| **6** (ACR-130) | 58.5 ± 1.6 | 35.3 ± 1.4 | 0 | 0 |
| **7** (ACR-147) | 54.7 ± 3.4 | 47.7 ± N/A | 0 | 0 |
| **8** (ACR-184) | 20.6 ± 3.7 | 61.1 ± 14.2 | Δ+260 (27%), Δ+520 (61%) | Δ+260 (14%) |
| **9** (ACR-195) | 57.9 ± 0.9 | 44.4 ± 2.0 | 0 | 0 |
| **10** (ACR-199) | 74.5 ± 3.0 | 64.6 ± 1.3 | 0 | 0 |
| **11** (ACR-300) | 32.3 ± 2.0 | 27.6 ± 0.4 | Δ+320 (100%) | 0 |
| **12** (ACR-338) | 55.2 ± 1.3 | 43.8 ± 2.3 | Δ+107 (40%), Δ+258 (18%), Δ+367 (14%) | 0 |
| **13** (ACR-345) | 48.0 ± 4.3 | 31.8 ± 0.1 | Δ+107 (39%) | 0 |

**Supporting Table S2.** X-Ray Crystal Diffraction Data Collection and Refinement Statistics

| | TEAD2-**14** (ACR-374) | TEAD2-**15** (ACR-378) | TEAD2-**16** (ACR-380) | TEAD3-**14** (ACR-374) |
|---|---|---|---|---|
| *Data collection* | | | | |
| Wavelength (Å) | 1.07216 | 1.00003 | 1.07216 | 0.99998 |
| Space group | C2 | C2 | C2 | P 21 21 21 |
| *Cell dimensions* | | | | |
| a, b, c (Å) | 122.39 61.45 79.53 | 121.62 61.72 79.41 | 121.67 61.22 79.52 | 65.27 121.76 153.22 |
| $\alpha, \beta, \gamma$ (°) | 90.00 117.69 90.00 | 90.00 117.71 90.00 | 90.00 117.72 90.00 | 90.00 90.00 90.00 |
| Resolution [a] (Å) | 54.19 – 2.60 (2.83 – 2.60) | 53.55 – 2.50 (2.80 – 2.50) | 57.62 – 2.23 (2.51 – 2.23) | 95.32 – 2.98 (3.29 – 2.98) |
| *Ellipsoidal Resolution* [b,c] (Å) | | | | |
| a* | 3.21 (0.993 a* - 0.117 c*) | 3.27 (0.998 a* + 0.063 c*) | 3.18 (0.999 a* - 0.041 c*) | 2.84 (a*) |
| b* | 2.69 (b*) | 2.78 (b*) | 2.70 (b*) | 4.23 (b*) |
| c* | 2.52 (- 0.430 a* + 0.903 c*) | 2.45 (- 0.635 a* + 0.772 c*) | 2.17 (- 0.535 a* + 0.845 c*) | 3.50 (c*) |
| $R_{sym}$ | 0.167 (0.948) | 0.133 (0.908) | 0.092 (0.581) | 0.301 (1.634) |
| $R_{meas}$ | 0.195 (1.102) | 0.157 (1.090) | 0.113 (0.680) | 0.314 (1.719) |
| $R_{pim}$ | 0.100 (0.560) | 0.081 (0.598) | 0.065 (0.497) | 0.088 (0.522) |
| Total reflections | 44803 (2305) | 39375 (1763) | 39451 (2512) | 177466 (7197) |
| No. unique reflections | 11910 (596) | 10768 (538) | 13453 (673) | 14367 (719) |
| CC1/2 | 0.989 (0.571) | 0.993 (0.545) | 0.995 (0.804) | 0.996 (0.582) |
| $I/\sigma(I)$ | 7.0 (1.3) | 7.8 (1.3) | 6.4 (1.4) | 9.0 (1.5) |
| Completeness spherical (%) | 73.4 (16.7) | 59.3 (10.5) | 53.0 (9.0) | 56.0 (11.2) |
| Completeness ellipsoidal [d] (%) | 91.0 (47.1) | 84.1 (50.8) | 88.4 (69.6) | 88.8 (75.3) |
| Multiplicity | 3.8 (3.9) | 3.7 (3.3) | 2.9 (3.7) | 12.4 (10.0) |
| Wilson B-factor | 41.52 | 40.21 | 33.47 | 45.08 |
| *Refinement* | | | | |
| Resolution (Å) | 53.46 – 2.60 (2.70 – 2.60) | 53.55 – 2.50 (2.59 – 2.50) | 57.62 – 2.23 (2.31 – 2.23) | 95.32 – 2.98 (3.21 – 2.98) |
| No. unique reflections | 11902 | 10762 | 13449 | 14360 |
| $R_{work}$ | 0.2083 | 0.2195 | 0.1952 | 0.2203 |
| $R_{free}$ | 0.2511 | 0.2682 | 0.2242 | 0.2815 |
| *R.m.s.d values* | | | | |
| Bond lengths (Å) | 0.004 | 0.003 | 0.003 | 0.003 |

| | | | | |
|---|---|---|---|---|
| Bond angles (°) | 0.891 | 0.622 | 0.756 | 0.687 |
| *No. atoms* | | | | |
| Protein | 3345 | 3333 | 3362 | 6723 |
| Ligand | 46 | 46 | 46 | 92 |
| solvent | 38 | 14 | 71 | 0 |
| *B-factors* (Å$^2$) | | | | |
| Protein | 46.50 | 44.22 | 39.50 | 47.56 |
| ligand | 77.47 | 48.52 | 65.31 | 47.50 |
| solvent | 34.00 | 34.44 | 36.67 | NA |
| *Ramachandran plot* | | | | |
| Favored (%) | 97.4 | 96.2 | 98.7 | 98.8 |
| Allowed (%) | 2.6 | 3.8 | 1.3 | 1.2 |
| Clashscore | 5.54 | 5.41 | 5.09 | 10.5 |
| Rotamer outliers (%) | 0.82 | 1.09 | 0.27 | 0.13 |
| *PDB code* | 8E0I | 8E0K | 8E0J | 8FKN |

(a) Highest-resolution shell values are shown in parentheses.
(b) Statistics are for data truncated using STARANISO server to remove reflections that were poorly measured due to the severe anisotropy presented in the data sets.
(c) Resolution limits for three directions in the reciprocal space.
(d) Anisotropic completeness was obtained by fitting an ellipsoid using least squares to the resulting cut-off surface points. Cut-off criteria were Rpim < 0.6, I/σ > 1.2 and CC1/2 > 0.5.

**Supporting Figure S8.** HPLC Data for Synthesized Acrylamides

Compound **1** (ACR-021) HPLC UV 220 nm, UV 254 nm

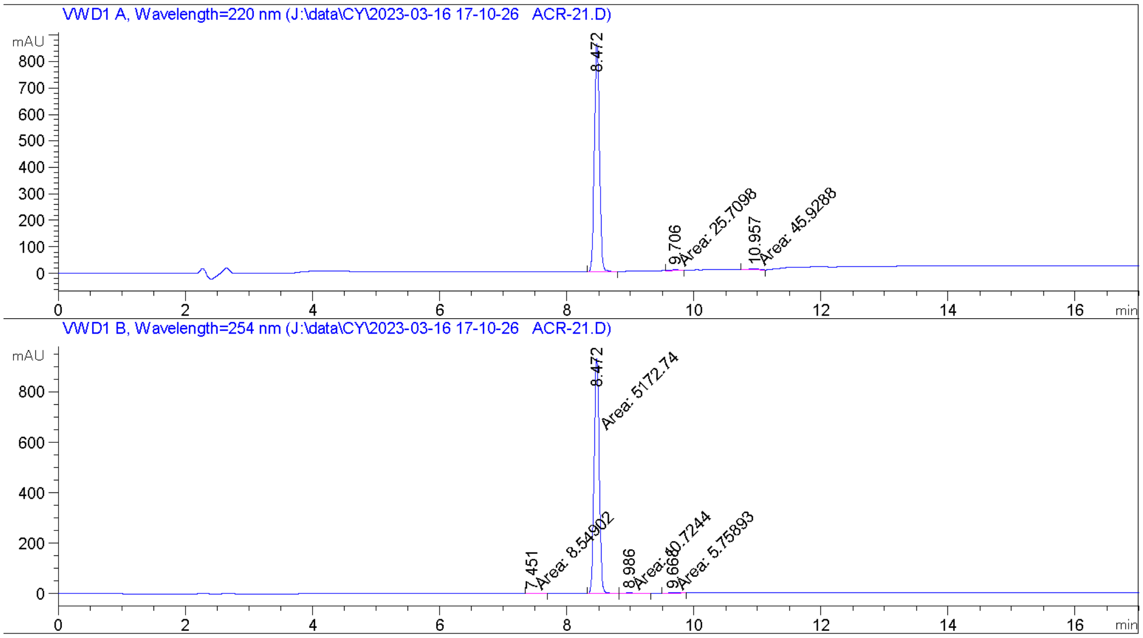


---

| | | |
|---|---|---|
| UV 220 nm: | **RT: 8.472 min** | **Area: 98.52 %** |
| | RT: 9.706 min | Area: 0.53 % |
| | RT: 10.957 min | Area: 0.95 % |
| UV 254 nm: | RT: 7.451 min | Area: 0.16 % |
| | **RT: 8.472 min** | **Area: 99.52 %** |
| | RT: 8.986 min | Area: 0.21 % |
| | RT: 9.668 min | Area: 0.11 % |

Compound **14** (ACR-374) HPLC UV 220 nm, UV 254 nm

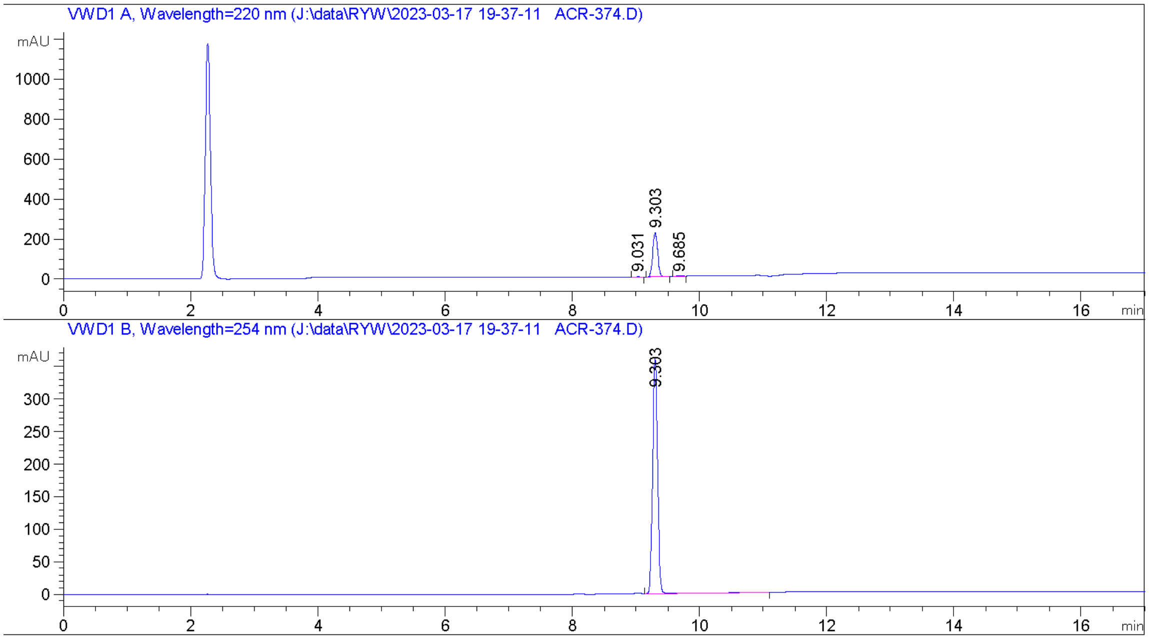


---

| | | |
|---|---|---|
| UV 220 nm: | RT: 9.031 min | Area: 1.02 % |
| | **RT: 9.303 min** | **Area: 97.89 %** |
| | RT: 9.685 min | Area: 1.08 % |
| UV 254 nm: | **RT: 9.303 min** | **Area: 100 %** |

---

Compound **15** (ACR-378) HPLC UV 220 nm, UV 254 nm

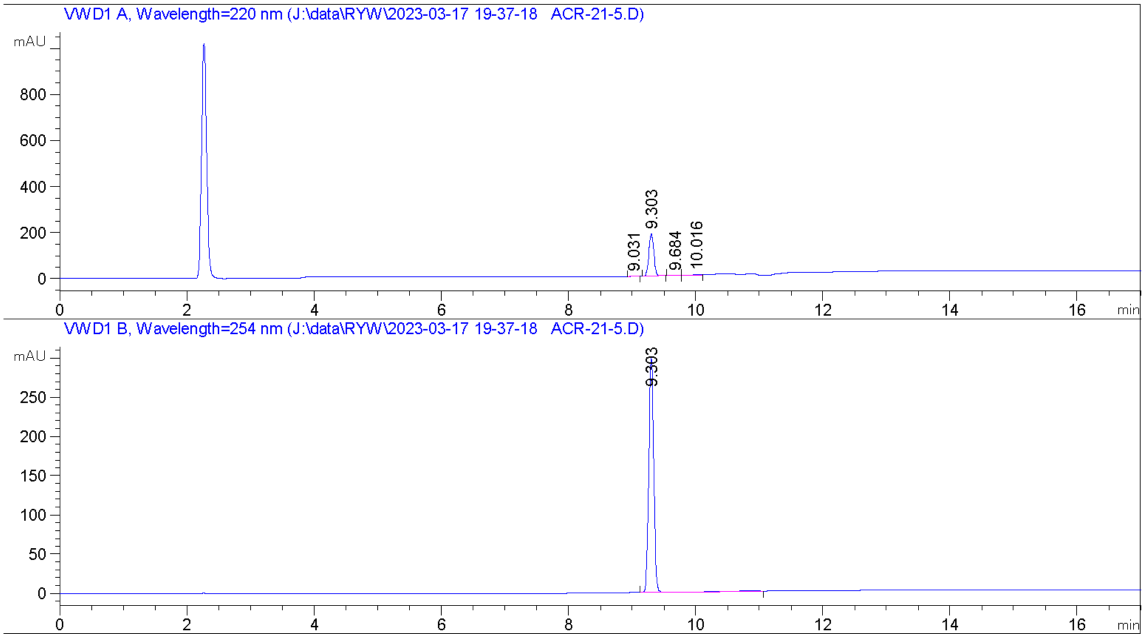


---

| | | |
|---|---|---|
| UV 220 nm: | RT: 9.031 min | Area: 0.95 % |
| | **RT: 9.303 min** | **Area: 96.86 %** |
| | RT: 9.684 min | Area: 2.19 % |
| UV 254 nm: | **RT: 9.303 min** | **Area: 100 %** |

---

Compound **16** (ACR-380) HPLC UV 220 nm, UV 254 nm

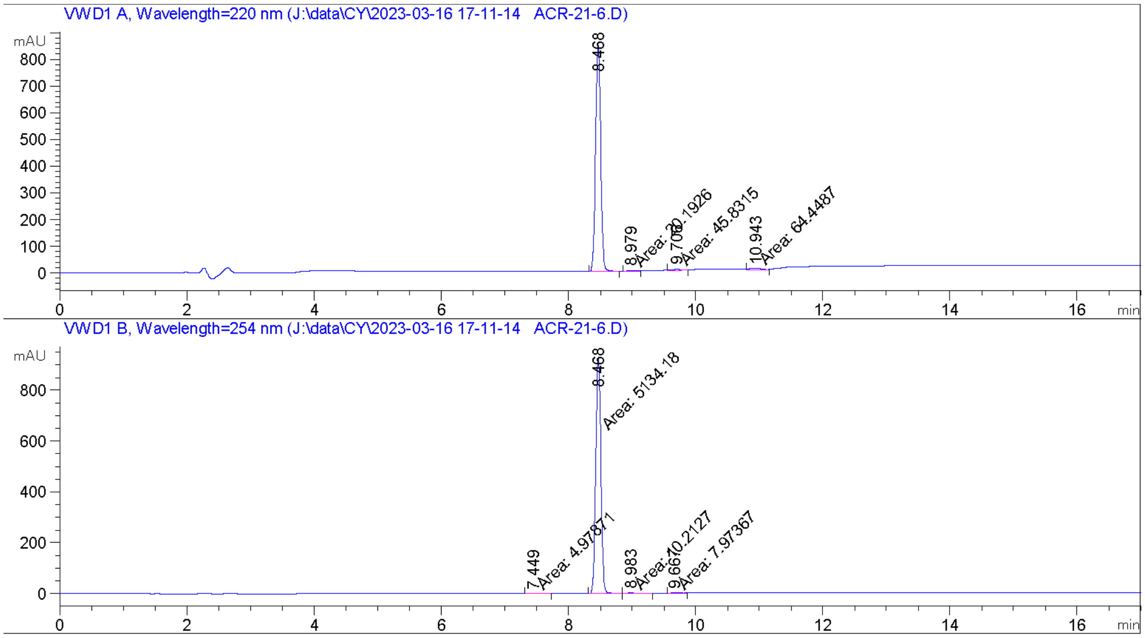


| | | |
|---|---|---|
| UV 220 nm: | **RT: 8.468 min** | **Area: 97.32 %** |
| | RT: 8.979 min | Area: 0.42 % |
| | RT: 9.706 min | Area: 0.94 % |
| | RT: 10.943 min | Area: 1.33 % |
| UV 254 nm: | RT: 7.449 min | Area: 0.10 % |
| | **RT: 8.468 min** | **Area: 99.55 %** |
| | RT: 8.983 min | Area: 0.20 % |
| | RT: 9.667 min | Area: 0.15 % |

Compound **17** (ACR-375) HPLC UV 220 nm, UV 254 nm

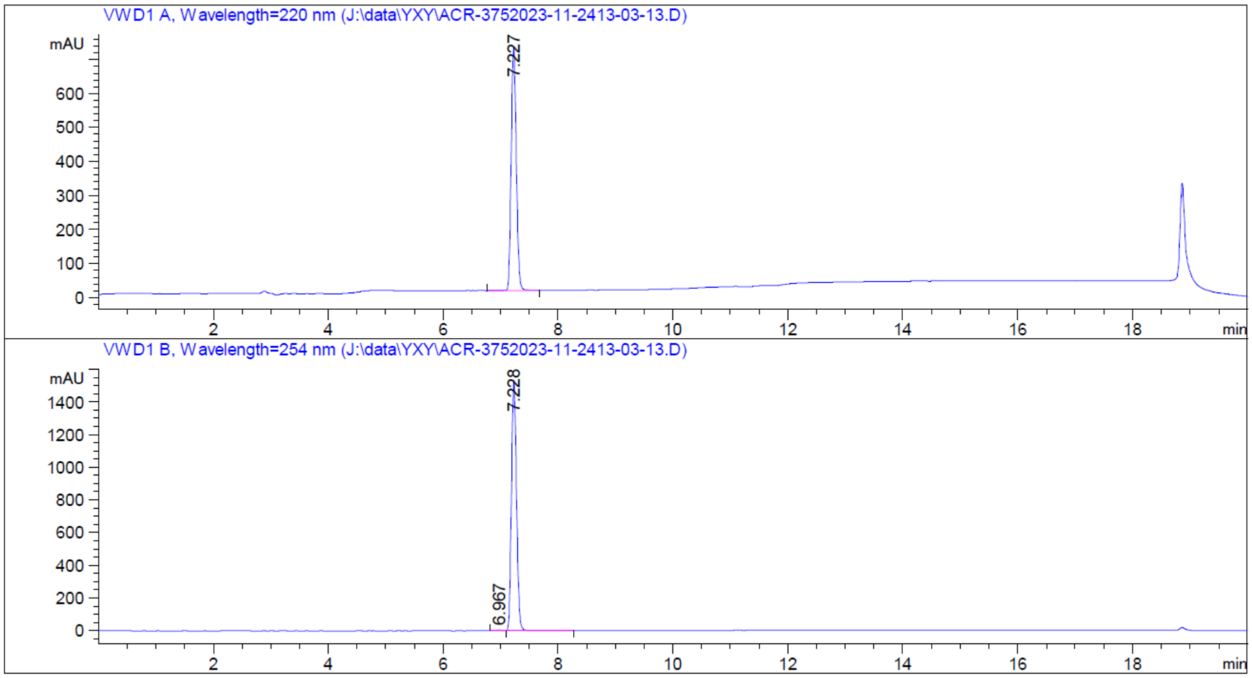


---

| | | |
|---|---|---|
| UV 220 nm: | **RT: 7.227 min** | **Area: 100 %** |
| UV 254 nm: | RT: 6.967 min | Area: 0.26 % |
| | **RT: 7.228 min** | **Area: 99.74 %** |

---

### Chemical Synthesis

Scheme 1. Synthesis of 1 (ACR-021)

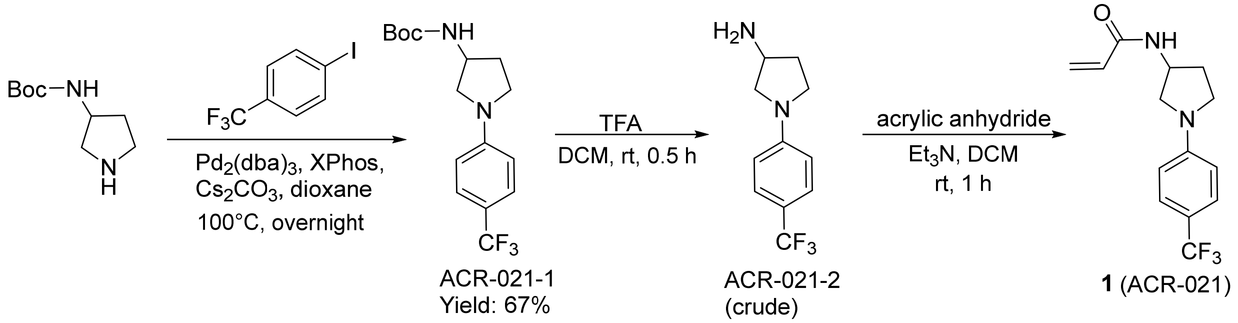


Synthesis of *tert*-butyl (1-(4-(trifluoromethyl)phenyl)pyrrolidin-3-yl)carbamate (ACR-021-1). A mixture of tert-butyl pyrrolidin-3-ylcarbamate (186.0 mg, 1.0 mmol), 1-iodo-4-(trifluoromethyl)benzene (326 mg, 1.2 mmol), $Pd_2(dba)_3$ (91.6 mg, 0.1 mmol), XPhos (95.2 mg, 0.2 mmol), and $Cs_2CO_3$ (650 mg, 2.0 mmol) in dioxane (30 mL) was stirred at 100 $^oC$ under $N_2$ overnight. The reaction mixture was cooled to rt and filtered over Celite. The filtrate was concentrated. The residue was dissolved in ethyl acetate (40 mL) and washed with brine. The solution was dried over $Na_2SO_4$. The solution was filtered, and the filtrate was concentrated. The residue was purified by column chromatography (PE/EA=10/1) to give *tert*-butyl (1-(4-(trifluoromethyl)phenyl)pyrrolidin-3-yl)carbamate (220 mg, 67% yield).

LRMS (m/z) for $C_{16}H_{22}F_3N_2O^+$ $[M+H]^+$: calculated 331.2, found 331.2.

Synthesis of 1-(4-(trifluoromethyl)phenyl)pyrrolidin-3-amine (ACR-021-2). The method was same as ACR-374-6 to give 1-(4-(trifluoromethyl)phenyl)pyrrolidin-3-amine (75 mg crude).

LRMS (m/z) for $C_{11}H_{14}F_3N_2^+$ $[M+H]^+$: calculated 231.1, found 231.1.

Synthesis of *N*-(1-(4-(trifluoromethyl)phenyl)pyrrolidin-3-yl)acrylamide [1 (ACR-021)]. The method was same as ACR-374 to give *N*-(1-(4-(trifluoromethyl)phenyl)pyrrolidin-3-yl)acrylamide.

LRMS (m/z) for $C_{14}H_{16}F_3N_2O^+$ $[M+H]^+$: calculated 285.1, found 285.1; $^1H$ NMR (400 MHz, $CDCl_3$): δ 7.46 (d, *J*=16.4 Hz, 2H), 6.56 (d, *J*=8.8 Hz, 2H), 6.34-6.30 (m, 1H), 6.11-6.04 (m, 1H), 5.84 (br, 1H), 5.69-5.66 (m, 1H), 4.72-4.69 (m, 1H), 3.68-3.63 (m, 1H), 3.52-3.46 (m, 1H), 3.42-3.38 (m, 1H), 3.37-3.24 (m, 1H), 2.38-2.31 (m, 1H), 2.01-2.03 (m, 1H).

Scheme 2. Synthesis of 14 (ACR-374)

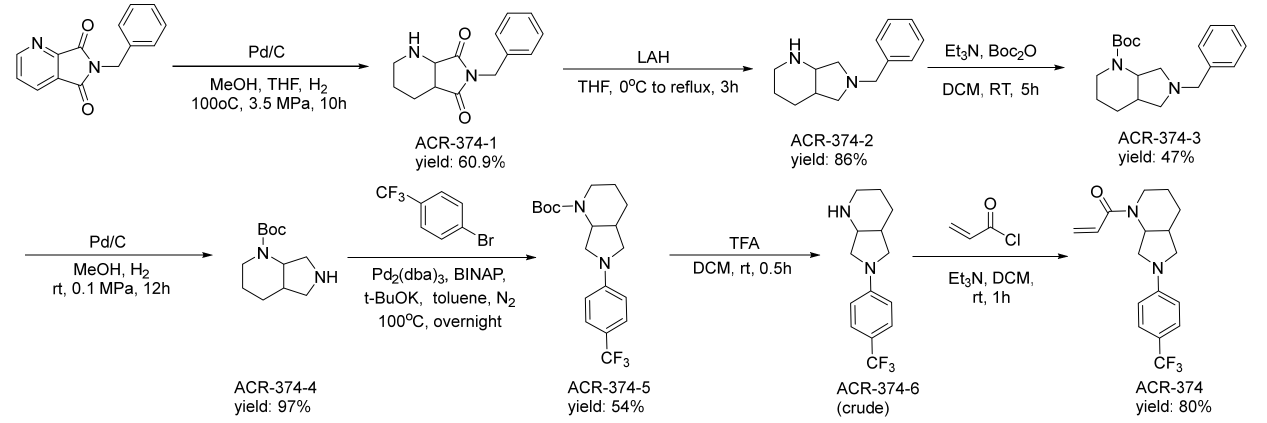


Synthesis of 6-benzylhexahydro-5*H*-pyrrolo[3,4-*b*]pyridine-5,7(6*H*)-dione (ACR-374-1). A mixture of 6-benzyl-5*H*-pyrrolo[3,4-*b*]pyridine-5,7(6*H*)-dione (4.0 g), 5% Pd/C (500 mg), and MeOH (10 mL) and THF (10 mL) was heated to 100 °C under hydrogen atmosphere in autoclave at a pressure of 3.5 MPa for 10 h. After cooling to rt, Pd/C was filtered out immediately, and the filtrate was concentrated to yield a light-yellow oil (2.5 g, 60.9% yield).

LRMS (m/z) for $C_{14}H_{16}N_2O_2^+$ $[M+H]^+$: calculated 245.3, found 245.3.

Synthesis of 6-benzyloctahydro-1*H*-pyrrolo[3,4-*b*]pyridine (ACR-374-2). To a solution of 6-benzylhexahydro-5H-pyrrolo[3,4-b]pyridine-5,7(6H)-dione (1.0 g, 4.09 mmol) in THF (40 mL) was added LAH (0.839 g, 20.4 mmol) in portions at 0 °C. Then the reaction was refluxed under $N_2$ for 3 h. The reaction was quenched by water and NaOH. The mixture was filtered, and the filtrate was concentrated. The residue was purified by silica gel chromatography (eluting with petroleum ether/EtOAc=10:1-1:2) to give 6-benzyloctahydro-1*H*-pyrrolo[3,4-*b*]pyridine (760 mg, 86% yield).

LRMS (m/z) for $C_{14}H_{21}N_2O_2{}^+$ $[M+H]^+$: calculated 217.3, found 217.3.

Synthesis of *tert*-butyl 6-benzyloctahydro-1H-pyrrolo[3,4-b]pyridine-1-carboxylate (ACR-374-3). To a solution of 6-benzyloctahydro-1*H*-pyrrolo[3,4-*b*]pyridine (760 mg, 3.5 mmol) and $Et_3N$ (1 mL, 3 mmol) in DCM was added $Boc_2O$ (6.6 g, 30.4 mmol). The mixture was stirred at room temperature for 5 h. The mixture was poured into water and the mixture was extracted with EA (100 mL x3). The organic phase was washed with water and brine. The solution was dried over $Na_2SO_4$ and filtered. The filtrate was concentrated and the residue was purified by column chromatography to give *tert*-butyl 6-benzyloctahydro-1*H*-pyrrolo[3,4-b]pyridine-1-carboxylate as a colorless oil (1.0 g, 47% yield).

LRMS (m/z) for $C_{19}H_{28}N_2O_2^+$ $[M+H]^+$: calculated 317.5, found 317.5.

Synthesis of *tert*-butyl octahydro-1*H*-pyrrolo[3,4-*b*]pyridine-1-carboxylate (ACR-374-4). A mixture of *tert*-butyl 6-benzyloctahydro-1*H*-pyrrolo[3,4-*b*]pyridine-1-carboxylate (400 mg), 5% Pd/C (100 mg) and MeOH (20 mL) was stirred at rt under hydrogen atmosphere in autoclave at a pressure of 0.1 MPa for 12 h. Pd/C was filtered out immediately, and the filtrate was concentrated to yield a light yellow oil (280 mg, 97% yield).

LRMS (m/z) for $C_{12}H_{22}N_2O_2^+$ $[M+H]^+$: calculated 227.3, found 227.3.

Synthesis of *tert*-butyl 6-(4-(trifluoromethyl)phenyl)octahydro-1*H*-pyrrolo[3,4-*b*]pyridine-1-carboxylate (ACR-374-5). A mixture of *tert*-butyl octahydro-1*H*-pyrrolo[3,4-*b*]pyridine-1-

carboxylate (226 mg, 1 mmol), 1-bromo-4-(trifluoromethyl)benzene (270 mg, 1.2 mmol), $Pd_2(dba)_3$ (91.6 mg, 0.1 mmol), BINAP (124.4 mg, 0.2 mmol), and *t*-BuOK (224 mg, 2.0 mmol) in toluene (30 mL) was stirred at 100 °C under $N_2$ overnight. The reaction mixture was cooled to rt and filtered over Celite. The filtrate was concentrated. The residue was dissolved in ethyl acetate (40 mL) and washed with brine. The solution was dried over $Na_2SO_4$. The solution was filtered, and the filtrate was concentrated. The residue was purified by column chromatography (PE/EA=8/1) to give *tert*-butyl 6-(4-(trifluoromethyl)phenyl)octahydro-1*H*-pyrrolo[3,4-*b*]pyridine-1-carboxylate (200 mg, 54% yield).

LRMS (m/z) for $C_{19}H_{25}F_3N_2O_2^+$ $[M+H]^+$: calculated 371.4, found 371.4; $^1H$ NMR (400 MHz, $CDCl_3$): δ 7.44 (d, *J*=8.4Hz, 1H), 6.52 (d, *J*=8.8Hz, 1H), 4.81 (m,1H), 4.07-4.05 (m,1H), 3.49-3.43 (m, 2H), 3.31-3.26 (m, 1H), 3.17-3.14 (m, 1H), 2.82-2.76 (m, 1H), 2.32-2.28 (m, 1H), 1.81-1.70 (m, 2H), 1.48 (s, 9H).

Synthesis of 6-(4-(trifluoromethyl)phenyl)octahydro-1*H*-pyrrolo[3,4-*b*]pyridine (ACR-374-6). *tert*-butyl 6-(4-(trifluoromethyl)phenyl)octahydro-1*H*-pyrrolo[3,4-*b*]pyridine-1-carboxylate (100 mg) was dissolved with DCM (2 mL), and $CF_3COOH$ (0.5 mL) was added. The mixture was stirred for 30 min at rt and concentrated. The residue was dissolved with $H_2O$ and washed with $Et_2O$. The water layer was basified with $NaHCO_3$, and the mixture was extracted with DCM. The combined organic layers were concentrated to afford the crude product.

LRMS (m/z) for $C_{14}H_{17}F_3N_2^+$ $[M+H]^+$: calculated 271.4, found 271.4.

Synthesis of 1-(6-(4-(trifluoromethyl)phenyl)octahydro-1*H*-pyrrolo[3,4-*b*]pyridin-1-yl)prop-2-en-1-one [14 (ACR-374)]. To a mixture of 6-(4-(trifluoromethyl)phenyl)octahydro-1*H*-pyrrolo[3,4-*b*]pyridine (80 mg) and $Et_3N$ (50 mg, 0.5 mmol) in DCM (10 mL) was added acryloyl chloride (8.6 mg, 0.095 mmol). The mixture was stirred at rt for 1 h. The mixture was quenched with water and extracted with DCM (20 mL x2). The combined organic phase was washed with water and brine, and dried over $Na_2SO_4$. The solvent was concentrated, and the residue was purified by preparative TLC (PE/EA = 1:1) to give the desired product as a white solid (76 mg, 80% yield).

LRMS (m/z) for $C_{17}H_{19}F_3N_2O^+$ $[M+H]^+$: calculated 325.3, found 325.3; $^1H$ NMR (400 MHz, $CDCl_3$): δ 7.46-7.44 (m, 2H), 6.66-6.51 (m, 3H), 6.34-6.29 (m, 1H), 5.74-5.71 (m, 1H), 5.35-5.25 (m, 0.5H), 4.76-4.63 (m, 0.5H), 3.94-3.85 (m, 1H), 3.53-3.49 (m, 3H), 3.22-3.19 (m, 2H), 2.39-2.36 (m, 1H), 1.89-1.79 (m, 2H); $^{13}C$ NMR (400 MHz, *$d_6$*-DMSO): δ 150.47, 129.23, 127.96, 127.25, 126.61, 126.57, 124.57, 115.69, 115.38, 111.45, 52.78, 25.97.

Scheme 3. Synthesis of 15 (ACR-378)

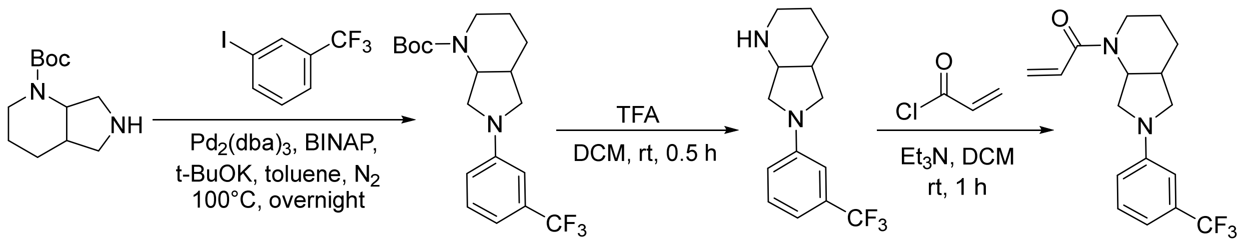


ACR-374-4

ACR-378-1
Yield: 97%

ACR-378-2
(crude)

**15** (ACR-378)
Yield: 34%

Synthesis of *tert*-butyl 6-(3-(trifluoromethyl)phenyl)octahydro-1*H*-pyrrolo[3,4-*b*]pyridine-1-carboxylate (ACR-378-1). The method was same as ACR-374-5 to give *tert*-butyl 6-(3-(trifluoromethyl)phenyl)octahydro-1*H*-pyrrolo[3,4-*b*]pyridine-1-carboxylate (160 mg, 97 % yield).

LRMS (m/z) for $C_{19}H_{25}F_3N_2O_2^+$ [M+H]$^+$: calculated 371.4, found 371.4.

Synthesis of 6-(3-(trifluoromethyl)phenyl)octahydro-1*H*-pyrrolo[3,4-*b*]pyridine (ACR-378-2). The method was same as ACR-374-6 to give 6-(3-(trifluoromethyl)phenyl)octahydro-1*H*-pyrrolo[3,4-*b*]pyridine (crude).

LRMS (m/z) for $C_{14}H_{17}F_3N_2^+$ [M+H]$^+$ : calculated 271.4, found 271.4.

Synthesis of 1-(6-(3-(trifluoromethyl)phenyl)octahydro-1*H*-pyrrolo[3,4-*b*]pyridin-1-yl)prop-2-en-1-one [15 (ACR-378)]. The method was same as ACR-374 to give 1-(6-(3-(trifluoromethyl)phenyl)octahydro-1H-pyrrolo[3,4-b]pyridin-1-yl)prop-2-en-1-one (47 mg, 34% yield).

LRMS (m/z) for $C_{17}H_{19}F_3N_2O^+$ [M+H]$^+$: calculated 325.3, found 325.3; $^1$H NMR (400 MHz, $CDCl_3$): δ 7.32-7.26 (m, 1H), 6.94 (s, 1H), 6.92-6.58 (m, 2H), 6.30 (d, *J*=16.8 Hz, 1H), 5.75 (d, *J*=10.8 Hz, 1H), 5.35 (m, 0.5H), 4.67 (m, 1H), 3.93 (m, 0.5H), 3.56-3.31 (m, 3H), 3.22-3.19 (m, 1.5H), 2.76-2.71 (m, 0.5H), 2.38-2.37 (m, 1H), 1.90-1.80 (m, 2H), 1.51-1.49 (m, 2H); $^{13}$C NMR (400 MHz, $CDCl_3$): δ 147.53, 129.58, 127. 81, 125.81, 123.10, 114.35, 107.70, 107.66, 52.64, 50.81, 45.39, 41.86, 35.69, 26.01.

Scheme 4. Synthesis of 16 (ACR-380)

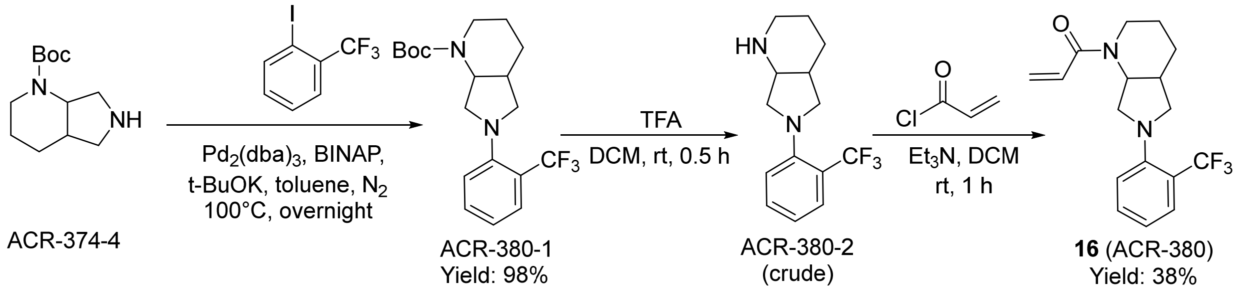


Synthesis of *tert*-butyl 6-(2-(trifluoromethyl)phenyl)octahydro-1*H*-pyrrolo[3,4-*b*]pyridine-1-carboxylate (ACR-380-1). The method was same as ACR-374-5 to give *tert*-butyl 6-(2-(trifluoromethyl)phenyl)octahydro-1*H*-pyrrolo[3,4-*b*]pyridine-1-carboxylate (180 mg, 98 % yield).

LRMS (m/z) for $C_{19}H_{25}F_3N_2O_2^+$ [M+H]$^+$: calculated 371.4, found 371.4.

Synthesis of 6-(2-(trifluoromethyl)phenyl)octahydro-1*H*-pyrrolo[3,4-*b*]pyridine (ACR-380-2). The method was same as ACR-374-6 to give 6-(2-(trifluoromethyl)phenyl)octahydro-1*H*-pyrrolo[3,4-*b*]pyridine (crude).

LRMS (m/z) for $C_{14}H_{17}F_3N_2^+$ [M+H]$^+$: calculated 271.4, found 271.4.

Synthesis of 1-(6-(2-(trifluoromethyl)phenyl)octahydro-1*H*-pyrrolo[3,4-*b*]pyridin-1-yl)prop-2-en-1-one [16 (ACR-380)]. The method was same as ACR-374 to give 1-(6-(2-(trifluoromethyl)phenyl)octahydro-1*H*-pyrrolo[3,4-*b*]pyridin-1-yl)prop-2-en-1-one (44.2 mg, 38 % yield).

LRMS (m/z) for $C_{17}H_{19}F_3N_2O^+$ $[M+H]^+$: calculated 325.3, found 325.3; $^1$H NMR (400 MHz, $CDCl_3$): δ 7.57 (d, *J*=8.0 Hz, 1H), 7.48 (t, *J*=8.0 Hz, 1H), 7.13 (d, *J*=7.6 Hz, 1H), 6.94 (t, *J*=7.6 Hz, 1H), 6.88-6.84 (m, 1H), 6.14-6.09 (m, 1H), 5.69-5.67 (m, 1H), 5.07 (m, 0.5H), 4.80 (m, 0.5H), 4.39 (m, 0.5H), 3.98 (m, 0.5H), 3.69-3.65 (m, 1H), 3.21-2.96 (m, 4H), 2.34 (m, 1H), 1.76-1.73 (m, 2H), 1.44-1.41 (m, 2H).

Scheme 5. Synthesis of 17 (ACR-375)

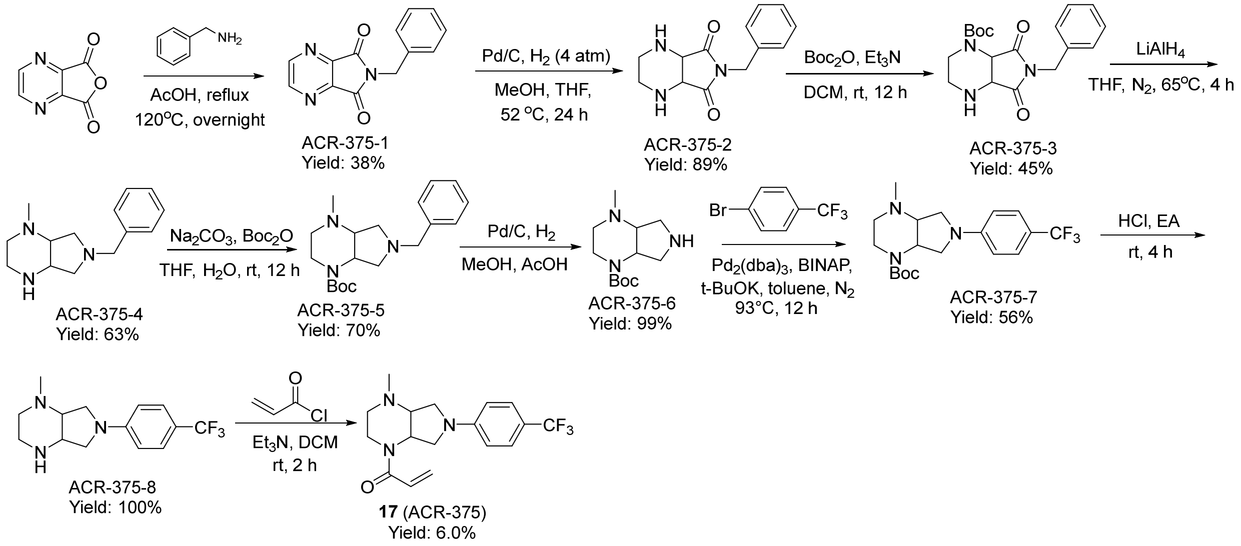


Synthesis of 6-benzyl-5*H*-pyrrolo[3,4-*b*]pyrazine-5,7(6*H*)-dione (ACR-375-1). To a solution of furo[3,4-*b*]pyrazine-5,7-dione (2.0 g, 13.3 mmol) in AcOH (20 mL) was added phenylmethanamine (1.43 g, 13.3 mmol) at rt. The mixture was stirred at 120 °C overnight. The solvent was removed under vacuum. The residue was purified by column chromatography (DCM/EA = 10:1) to give 6-benzyl-5*H*-pyrrolo[3,4-*b*]pyrazine-5,7(6*H*)-dione (1.2 g, 38% yield).

LCMS (m/z) for $C_{13}H_{10}N_3O_2$ $[M+H]^+$ : calculated 239.1, found 254.0.

Synthesis of 6-benzyltetrahydro-1*H*-pyrrolo[3,4-*b*]pyrazine-5,7(6*H*,7*aH*)-dione (ACR-375-2). A solution of 6-benzyl-5*H*-pyrrolo[3,4-*b*]pyrazine-5,7(6*H*)-dione (1.2 g, 2.1 mmol) and palladium on carbon (2.0 g, 10% wt) in MeOH/THF (1:1, v/v, 140 mL) was stirred at 52 °C under $H_2$ (4 atm) for 24 h. The mixture was filtered and concentrated to give crude 6-benzyltetrahydro-1*H*-pyrrolo[3,4-*b*]pyrazine-5,7(6*H*,7*aH*)-dione (1.1 g, 89.4% yield).

LRMS (m/z) for $C_{13}H_{15}N_3O_2$$[M+H]^+$: calculated 246.3, found 246.2.

Synthesis of *tert*-butyl 6-benzyl-5,7-dioxooctahydro-1*H*-pyrrolo[3,4-*b*]pyrazine-1-carboxylate (ACR-375-3). To a mixture of 6-benzyltetrahydro-1*H*-pyrrolo[3,4-b]pyrazine-5,7(6*H*,7*aH*)-dione (1.1 g, 4.49 mmol) and $Et_3N$ (1.36 g, 13.46 mmol) in DCM (50 mL) was added $Boc_2O$ (930.6 mg, 4.26 mmol) at 0 °C. The mixture was stirred at rt for 12 h. The mixture was quenched with water and extracted with DCM (20 mL x2). The combined organic phase was washed with water and brine, and dried over $Na_2SO_4$. The solvent was concentrated, and the residue was purified by column chromatography on silica gel (DCM/EA = 10:1) to give the desired product as a white solid (700 mg, 45% yield).

LRMS (m/z) for $C_{18}H_{24}N_3O_4^+$ $[M+H]^+$: calculated 346.4 found 346.4.

Synthesis of 6-benzyl-1-methyloctahydro-1*H*-pyrrolo[3,4-*b*]pyrazine (ACR-375-4). To a mixture of *tert*-butyl 6-benzyl-5,7-dioxooctahydro-1*H*-pyrrolo[3,4-*b*]pyrazine-1-carboxylate (700 mg, 2.02 mmol) in THF was added $LiAlH_4$ (698.3 mg, 18.37 mmol). The mixture was stirred at 65 $^{o}C$ for 4 h under the atmosphere of $N_2$. The mixture was quenched with MeOH/$H_2O$ (5 mL/5 mL) at 0 $^{o}C$. The mixture was added onto anhydrous $Na_2SO_4$ and filtered. The filtrate was concentrated to give 6-benzyl-1-methyloctahydro-1*H*-pyrrolo[3,4-*b*]pyrazine (1.12 g, 63% yield).

LRMS (m/z) for $C_{14}H_{22}N_3^+$ $[M+H]^+$: calculated 232.3, found 232.3.

Synthesis of *tert*-butyl 6-benzyl-4-methyloctahydro-1*H*-pyrrolo[3,4-b]pyrazine-1-carboxylate (ACR-375-5). To a mixture of 6-benzyl-1-methyloctahydro-1*H*-pyrrolo[3,4-*b*]pyrazine (500 mg, 2.16 mmol) in THF/$H_2O$ (10 mL/10 mL) were added $Boc_2O$ (943.7 mg, 4.33 mmol) and $Na_2CO_3$ (458.8 mg, 4.33 mmol). The mixture was stirred at rt for 12 h. The mixture was diluted with EA and washed with brine. The organic layer was dried over $Na_2SO_4$. The mixture was filtered, and the filtrate was concentrated. The residue was purified by column chromatography (PE/EA=3/1) to give *tert*-butyl 6-benzyl-4-methyloctahydro-1*H*-pyrrolo[3,4-*b*]pyrazine-1-carboxylate (500 mg, 70% yield).

LRMS (m/z) for $C_{19}H_{30}N_3O_2^+$ $[M+H]^+$: calculated 332.4, found 332.4.

Synthesis of *tert*-butyl 4-methyloctahydro-1*H*-pyrrolo[3,4-*b*]pyrazine-1-carboxylate (ACR-375-6). To a solution of *tert*-butyl 6-benzyl-4-methyloctahydro-1*H*-pyrrolo[3,4-b]pyrazine-1-carboxylate in MeOH (20 mL) were added Pd/C (1.2 g) and AcOH (3 drops). The mixture was stirred overnight at room temperature under the atmosphere of $H_2$. After completion of the reaction, the mixture was filtered over celite, and the filtrate was concentrated under vacuum. The residue was used for next step without further purification to give *tert*-butyl 4-methyloctahydro-1*H*-pyrrolo[3,4-*b*]pyrazine-1-carboxylate (360 mg, 98.9% yield).

LRMS (m/z) for $C_{12}H_{24}N_3O_2^+$ $[M+H]^+$: calculated 242.3, found 242.3.

Synthesis of *tert*-butyl 4-methyl-6-(4-(trifluoromethyl)phenyl)octahydro-1H-pyrrolo[3,4-b]pyrazine-1-carboxylate (ACR-375-7). To a solution of *tert*-butyl 4-methyloctahydro-1*H*-pyrrolo[3,4-*b*]pyrazine-1-carboxylate (200 mg, 0.83 mmol) in dry toluene (10 mL) were added 1-bromo-4-(trifluoromethyl)benzene (270.8 mg, 0.99 mmol), $Pd_2(dba)_3$ (76 mg, 0.083 mmol), BINAP (103.3 mg, 0.16 mmol) and *t*-BuOK (185.8 mg, 1.66 mmol). The mixture was stirred at 93 $^{o}C$ under the atmosphere of $N_2$ for 12 h. The mixture was diluted with EA and washed with brine. The organic layer was concentrated. The residue was purified by preparative TLC (PE/EA= 5:1) to give the desired product (180 mg, 56% yield).

LRMS (m/z) for $C_{19}H_{27}F_3N_3O_2^+$ $[M+H]^+$: calculated 386.2, found 386.2.

Synthesis of 1-methyl-6-(4-(trifluoromethyl)phenyl)octahydro-1*H*-pyrrolo[3,4-*b*]pyrazine (ACR-375-8). To a solution of *tert*-butyl 4-methyl-6-(4-(trifluoromethyl)phenyl)octahydro-1*H*-pyrrolo[3,4-*b*]pyrazine-1-carboxylate (180 mg, 0.47 mmol) in EA (6 mL) was added HCl/EA (4N, 3 mL), and the resulting mixture was stirred at rt for 4 h. After the completion of the reaction, the solvent was removed under vacuum to afford the desired product (133.2 mg, 100% yield).

LRMS (m/z) for $C_{14}H_{19}F_3N_3^+$ $[M+H]^+$: calculated 286.1, found 286.1.

Synthesis of 1-(4-methyl-6-(4-(trifluoromethyl)phenyl)octahydro-1*H*-pyrrolo[3,4-*b*]pyrazin-1-yl)prop-2-en-1-one [17 (ACR-375)]. To a solution of 1-methyl-6-(4-

(trifluoromethyl)phenyl)octahydro-1*H*-pyrrolo[3,4-*b*]pyrazine (133.2 mg, 0.47 mmol) in DCM (5 mL) was added $Et_3N$ (2 mL). Then acryloyl chloride (50.77 mg, 0.56 mmol) was added at 0 $^oC$. The mixture was warmed up to rt and stirred for 2 h. Then the mixture was diluted with EA and washed with brine. The organic layer was dried over $Na_2SO_4$. The mixture was filtered, and the filtrate was concentrated. The residue was purified by preparative HPLC to give the desired product (9.7 mg, 6.0% yield).

LRMS (m/z) for $C_{17}H_{20}F_3N_3O^+$ $[M+H]^+$: calculated 340.2, found 340.2; $^1H$ NMR (400 MHz, $CDCl_3$): δ 7.47 (d, *J*=8.6 Hz, 2H), 6.56 (q, *J*=9.5 Hz, 3H), 6.47-6.37 (m, 1H), 5.87 (dd, *J*=10.4, 1.6 Hz, 1H), 3.96 (d, *J*=12.0 Hz, 2H), 3.84 (s, 4H), 3.77-3.70 (m, 2H), 3.66 (s, 2H), 3.55 (d, *J*=12.4 Hz, 1H), 3.36 (t, *J*=5.2 Hz, 1H), 2.85 (s, 3H), 2.78 (s, 1H).

### REFERENCES


[1] K. Bum-Erdene, D. Zhou, G. Gonzalez-Gutierrez, M. K. Ghozayel, Y. Si, D. Xu, H. E. Shannon, B. J. Bailey, T. W. Corson, K. E. Pollok, C. D. Wells, S. O. Meroueh, *Cell Chem Biol* **2019**, *26*, 378-389 e313.

[2] K. Bum-Erdene, I. J. Yeh, G. Gonzalez-Gutierrez, M. K. Ghozayel, K. Pollok, S. O. Meroueh, *J Med Chem* **2023**, *66*, 266-284.

[3] A. Kaneda, T. Seike, T. Danjo, T. Nakajima, N. Otsubo, D. Yamaguchi, Y. Tsuji, K. Hamaguchi, M. Yasunaga, Y. Nishiya, M. Suzuki, J. I. Saito, R. Yatsunami, S. Nakamura, Y. Sekido, K. Mori, *Am J Cancer Res* **2020**, *10*, 4399-4415.

[4] H. Karatas, M. Akbarzadeh, H. Adihou, G. Hahne, A. V. Pobbati, E. Yihui Ng, S. M. Guéret, S. Sievers, A. Pahl, M. Metz, S. Zinken, L. Dötsch, C. Nowak, S. Thavam, A. Friese, C. Kang, W. Hong, H. Waldmann, *Journal of Medicinal Chemistry* **2020**, *63*, 11972-11989.

[5] W. Lu, J. Wang, Y. Li, H. Tao, H. Xiong, F. Lian, J. Gao, H. Ma, T. Lu, D. Zhang, X. Ye, H. Ding, L. Yue, Y. Zhang, H. Tang, N. Zhang, Y. Yang, H. Jiang, K. Chen, B. Zhou, C. Luo, *European Journal of Medicinal Chemistry* **2019**, *184*, 111767.

[6] T. Lu, Y. Li, W. Lu, T. Spitters, X. Fang, J. Wang, S. Cai, J. Gao, Y. Zhou, Z. Duan, H. Xiong, L. Liu, Q. Li, H. Jiang, K. Chen, H. Zhou, H. Lin, H. Feng, B. Zhou, C. L. Antos, C. Luo, *Acta Pharmaceutica Sinica B* **2021**, *11*, 3206-3219.

[7] aM. Fan, W. Lu, J. Che, N. P. Kwiatkowski, Y. Gao, H.-S. Seo, S. B. Ficarro, P. C. Gokhale, Y. Liu, E. A. Geffken, J. Lakhani, K. Song, M. Kuljanin, W. Ji, J. Jiang, Z. He, J. Tse, A. S. Boghossian, M. G. Rees, M. M. Ronan, J. A. Roth, J. D. Mancias, J. A. Marto, S. Dhe-Paganon, T. Zhang, N. S. Gray, *eLife* **2022**, *11*, e78810; bK. J. Kurppa, Y. Liu, C. To, T. Zhang, M. Fan, A. Vajdi, E. H. Knelson, Y. Xie, K. Lim, P. Cejas, A. Portell, P. H. Lizotte, S. B. Ficarro, S. Li, T. Chen, H. M. Haikala, H. Wang, M. Bahcall, Y. Gao, S. Shalhout, S. Boettcher, B. H. Shin, T. Thai, M. K. Wilkens, M. L. Tillgren, M. Mushajiang, M. Xu, J. Choi, A. A. Bertram, B. L. Ebert, R. Beroukhim, P. Bandopadhayay, M. M. Awad, P. C. Gokhale, P. T. Kirschmeier, J. A. Marto, F. D. Camargo, R. Haq, C. P. Paweletz, K.-K. Wong, D. A. Barbie, H. W. Long, N. S. Gray, P. A. Jänne, *Cancer Cell* **2020**, *37*, 104-122.e112.

[8] W. Lu, M. Fan, W. Ji, J. Tse, I. You, S. B. Ficarro, I. Tavares, J. Che, A. Y. Kim, X. Zhu, A. Boghossian, M. G. Rees, M. M. Ronan, J. A. Roth, S. M. Hinshaw, B. Nabet, S. M. Corsello, N. Kwiatkowski, J. A. Marto, T. Zhang, N. S. Gray, *Journal of Medicinal Chemistry* **2023**, *66*, 4617-4632.

[9] K. Nutsch, L. Song, E. Chen, M. Hull, A. K. Chatterjee, J. J. Chen, M. J. Bollong, *RSC Chem Biol* **2023**, *4*, 894-905.

[10] H. Hillen, A. Candi, B. Vanderhoydonck, W. Kowalczyk, L. Sansores-Garcia, E. C. Kesikiadou, L. Van Huffel, L. Spiessens, M. Nijs, E. Soons, W. Haeck, H. Klaassen, W. Smets, S. A. Spieser, A. Marchand, P. Chaltin, F. Ciesielski, F. Debaene, L. Chen, A. Kamal, S. L. Gwaltney, M. Versele, G. A. Halder, *Mol Cancer Ther* **2023**.

[11] A. V. Pobbati, X. Han, A. W. Hung, S. Weiguang, N. Huda, G. Y. Chen, C. Kang, C. S. Chia, X. Luo, W. Hong, A. Poulsen, *Structure* **2015**, *23*, 2076-2086.

[12] J. K. Holden, J. J. Crawford, C. L. Noland, S. Schmidt, J. R. Zbieg, J. A. Lacap, R. Zang, G. M. Miller, Y. Zhang, P. Beroza, R. Reja, W. Lee, J. Y. K. Tom, R. Fong, M. Steffek, S. Clausen, T. J. Hagenbeek, T. Hu, Z. Zhou, H. C. Shen, C. N. Cunningham, *Cell Reports* **2020**, *31*, 107809.

[13] aQ. Li, Y. Sun, G. K. Jarugumilli, S. Liu, K. Dang, J. L. Cotton, J. Xiol, P. Y. Chan, M. DeRan, L. Ma, R. Li, L. J. Zhu, J. H. Li, A. B. Leiter, Y. T. Ip, F. D. Camargo, X. Luo, R. L. Johnson, X. Wu, J. Mao, *Cell Stem Cell* **2020**, *26*, 675-692.e678; bY. Sun, L. Hu, Z. Tao, G. K. Jarugumilli, H. Erb, A. Singh, Q. Li, J. L. Cotton, P. Greninger, R. K. Egan, Y. Tony Ip, C. H. Benes, J. Che, J. Mao, X. Wu, *Nature Communications* **2022**, *13*, 6744.

[14] L. Li, R. Li, Y. Wang, *Bioorganic Chemistry* **2022**, *122*, 105707.

[15] T. T. Tang, A. W. Konradi, Y. Feng, X. Peng, M. Ma, J. Li, F.-X. Yu, K.-L. Guan, L. Post, *Molecular Cancer Therapeutics* **2021**, *20*, 986-998.

[16] L. Hu, Y. Sun, S. Liu, H. Erb, A. Singh, J. Mao, X. Luo, X. Wu, *eLife* **2022**, *11*, e80210.

[17] T. J. Hagenbeek, J. R. Zbieg, M. Hafner, R. Mroue, J. A. Lacap, N. M. Sodir, C. L. Noland, S. Afghani, A. Kishore, K. P. Bhat, X. Yao, S. Schmidt, S. Clausen, M. Steffek, W. Lee, P. Beroza, S. Martin, E. Lin, R. Fong, P. Di Lello, M. H. Kubala, M. N. Y. Yang, J. T. Lau, E. Chan, A. Arrazate, L. An, E. Levy, M. N. Lorenzo, H.-J. Lee, T. H. Pham, Z. Modrusan, R. Zang, Y.-C. Chen, M. Kabza, M. Ahmed, J. Li, M. T. Chang, D. Maddalo, M. Evangelista, X. Ye, J. J. Crawford, A. Dey, *Nature Cancer* **2023**, *4*, 812-828.

[18] T. Heinrich, C. Peterson, R. Schneider, S. Garg, D. Schwarz, J. Gunera, A. Seshire, L. Kötzner, S. Schlesiger, D. Musil, H. Schilke, B. Doerfel, P. Diehl, P. Böpple, A. R. Lemos, P. M. F. Sousa, F. Freire, T. M. Bandeiras, E. Carswell, N. Pearson, S. Sirohi, M. Hooker, E. Trivier, R. Broome, A. Balsiger, A. Crowden, C. Dillon, D. Wienke, *Journal of Medicinal Chemistry* **2022**, *65*, 9206-9229.
